# Chloride Homeostasis Regulates cGAS-STING Signaling

**DOI:** 10.1101/2024.04.08.588475

**Authors:** Jared Morse, Danna Wang, Serena Mei, Danielle Whitham, Colby Hladun, Costel C. Darie, Herman O. Sintim, Modi Wang, KaHo Leung

## Abstract

The cGAS–STING signaling pathway has emerged as a key mediator of inflammation. However, the roles of chloride homeostasis on this pathway are unclear. Here, we uncovered a correlation between chloride homeostasis and cGAS-STING signaling. We found that dysregulation of chloride homeostasis attenuates cGAS-STING signaling in a lysosome-independent manner. Treating immune cells with chloride channel inhibitors attenuated 2’3’-cGAMP production by cGAS and also suppressed STING polymerization, leading to reduced cytokine production. We also demonstrate that non-selective chloride channel blockers can suppress the NPC1 deficiency-induced, hyper-activated STING signaling in skin fibroblasts derived from Niemann Pick disease type C (NPC) patients. Our findings reveal that chloride homeostasis majorly affects cGAS-STING pathway and suggest a provocative strategy to dampen STING-mediated inflammation via targeting chloride channels.

**Highlights:**

- Chloride dysregulation attenuates cGAS-STING signaling in a lysosome-independent manner.
- Chloride dysregulation attenuates intracellular 2’3’-cGAMP production.
- Chloride dysregulation inhibits STING polymerization and STING-to-IRF3 signaling.
- Chloride channel blockers suppress NPC1 deficiency-induced, hyper-activated STING signaling.

**Graphical abstract:** 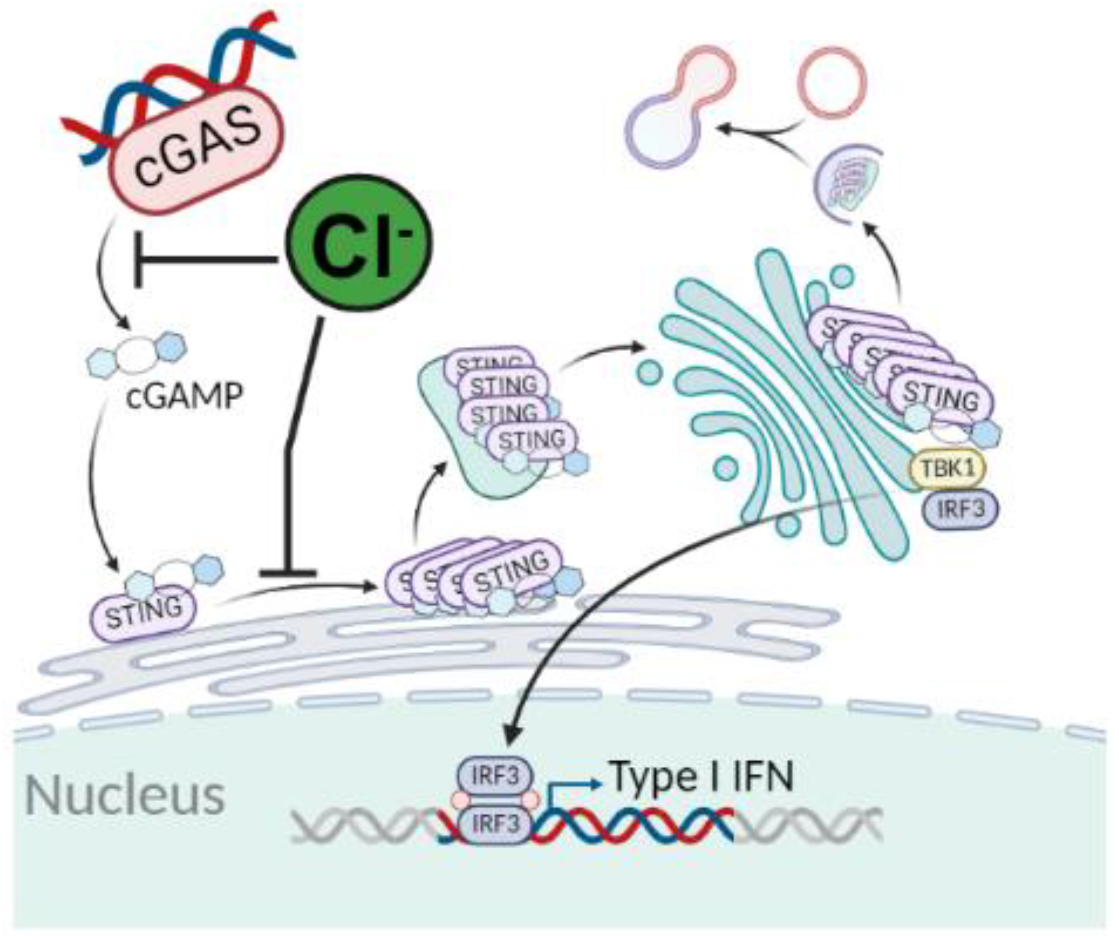

## Introduction

Within the realm of innate immune network, the cyclic GMP–AMP synthase (cGAS)–stimulator of interferon genes (STING) pathway plays a key role in recognizing and defending against foreign DNA entities.^1–5^ Increasing evidence however indicates that the dysregulation of the cGAS-STING pathway contributes to or accentuates the severity of many diseases.^6–8^ cGAS–STING signaling pathway is also considered as a crucial driver of chronic inflammation,^9–13^ which plays a critical role in neurodegenerative diseases,^13,14^ including Alzheimer’s disease,^15,16^ amyotrophic lateral sclerosis, Parkinson’s disease.^17^ Hence, the cGAS–STING signaling pathway has emerged as a pivotal mediator of inflammation^13^ and manipulating this signaling pathway is an emerging approach for development of therapeutic treatments.

Cytosolic DNA could be a result of pathogen infection or cellular stress. The cytosolic DNA sensor cGAS binds to double-stranded DNA (dsDNA) and synthesizes the cyclic dinucleotide 2′3′ cyclic GMP– AMP (2’3’-cGAMP) as the secondary messenger,^18–32^ which binds to STING on the endoplasmic reticulum membrane.^33,34^ Upon the binding of 2’3’-cGAMP, STING is activated and undergoes polymerization.^35^ It is subsequently translocated to Golgi via ERGIC (ER-Golgi intermediate complex) transport cluster.^36–39^ Upon translocation, STING activates TANK-binding kinase 1 (TBK1) and also brings transcription factor-interferon regulatory factor 3 (IRF3) in close proximity to TKB1, thereby facilitating IRF3 phosphorylation by TBK1.^40^ Subsequently, phospho-IRF3 dimerizes and locates to the nucleus where it regulates the transcription of type 1 interferons, such as IFN-β.^41–44^ The signaling is ultimately quenched via the degradation of the activated complex through autophagy.^45^ Beyond the regulation of interferons and cytokines, the cGAS-STING pathway has been found to affect cellular physiology via interfacing with other processes such as metabolism,^46–48^ cell death,^49,50^ and senescence.^51,52^ Meanwhile, these biological processes are mediated by chloride homeostasis.^53–61^

As the most abundant anion in the cells, the roles of chloride are beginning to come into focus. For example, chloride plays an important role in the regulation of cytoplasm composition,^62^ organelle volume,^63^ membrane potential,^64^ and cell excitability.^65^ However, how chloride homeostasis dysregulation affects the cGAS-STING pathway is yet to be elucidated. It has been reported that lysosomal chloride is crucial for lysosome function.^66–68^ Lysosomal chloride mediates the lysosomal calcium storage and directly regulates several key lysosomal enzymes.^59,69^ Dysregulation of lysosomal chloride reduces the context of autophagy,^70^ which is the process to quench the activated cGAS-STING signal. Furthermore, the proton channel property of STING has been recently uncovered. Activated STING polymer complexes can serve as a proton channel, inducing a pH increase in the Golgi.^71^ Chloride being the most abundant anion in the cell, plays a crucial role in functioning as the necessary counter ion during proton pumping, for example, maintaining Golgi membrane potential close to zero.^69,72–74^ Therefore, we hypothesize that chloride homeostasis is critical to cGAS-STING pathway. Dysregulation of chloride homeostasis or organellar chloride will consequently modulate cGAS-STING signaling.

Here, we investigated the influence of chloride homeostasis on cGAS-STING pathway by examining the dsDNA- and STING ligand-induced cGAS-STING signaling during chloride homeostasis dysregulation. We demonstrated that dysregulation of chloride homeostasis attenuates cGAS-STING signaling in a lysosome-independent manner. Treating immune cells with chloride channel inhibitors attenuated 2’3’-cGAMP production by cGAS and also suppressed STING polymerization, leading to reduced cytokine production. We then leveraged the non-selective chloride channel blockers to effectively suppress the hyper-activated STING signaling in skin fibroblasts derived from Niemann Pick disease type C (NPC) patients. These findings unveil a correlation between chloride homeostasis and cGAS-STING pathway, suggesting that chloride channel target therapy could be a potential strategy to suppress the pathology of NPC or STING associated diseases in which immunomodulation of the type I IFN pathway might show therapeutic benefit.

## Results

### Dysregulation of chloride homeostasis attenuates cytosolic DNA-stimulated cGAS-STING pathway

We initially conducted a functional investigation to elucidate the role of chloride homeostasis in cGAS-STING pathway. The activation of cGAS-STING pathway can be monitored by using an engineered IRF3 reporter cell THP1-BlueTM ISG. The activation of IRF3 induces the production of secreted reporter enzyme, secreted embryonic alkaline phosphatase (SEAP). The reporter enzyme is secreted into cell culture supernatant and IRF3 activation can be determined by reporter activity of SEAP (Figure 1a). To dysregulate the chloride homeostasis, we treated the reporter cells with non-selective chloride channel blockers such as NPPB,^75^ DIDS,^76,77^ IAA-94,^78–80^ and FFA (Supplementary Figure 1).^81^ We then transfected the cells with herring testes-DNA (HT-DNA) to stimulate the cGAS-STING pathway.^82–84^ When cells were pretreated with chloride channel blockers, the HT-DNA stimulated cGAS-STING signaling was inhibited in a dose dependent manner (Figure 1b-e). To assess whether this trend is consistently observed across various mammalian cells, we employed the murine sourced IRF3 reporter cells (RAW-Dual), allowing cGAS-STING pathway insight in murine macrophages. A similar inhibitory pattern was observed in RAW macrophages. Upon treatment of non-selective chloride channel blockers, the HT-DNA stimulated cGAS-STING signaling was attenuated (Figure 1f). To understand the impact of non-selective chloride channel blockers on the kinetics of signaling, we conducted a time-dependent analysis of the activation process. The temporal analysis showed that from 1-24h, chloride homeostasis dysregulation dynamically inhibits HT-DNA stimulated cGAS-STING signaling (Figure 1g). To eliminate the concern that non-selective chloride channel blockers reduce the signal of reporter cells by inducing cell death, we investigated the cytotoxicity of those chloride channel blockers. Cell viability remained unchanged upon the incubation with chloride channel blockers at the concentrations employed (Supplementary Figure 2 a-b).

**Figure 1.**
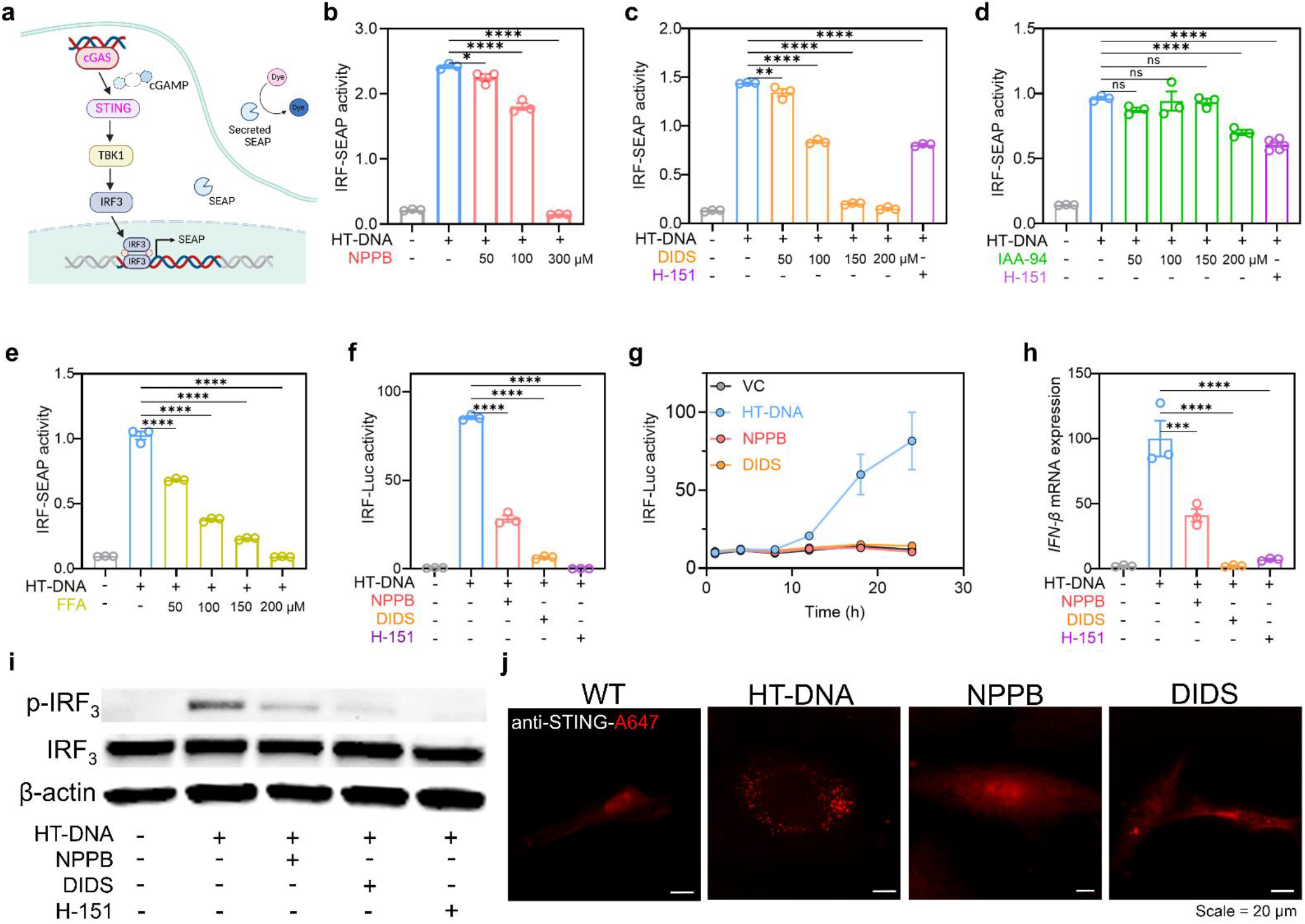
Dysregulation of cellular chloride attenuates cytosolic DNA-stimulated cGAS-STING pathway. (**a**) Schematic diagram illustrating the mechanism for monitoring cGAS-STING pathway activation via IRF3 reporter cells. Activation of IRF3 induces the production of secreted reporter enzyme. The reporter enzyme is secreted into cell culture supernatant and IRF_3_ activation can be determined by reporter activity using (**b**−**e**) colorimetric (OD at 655nm) or (**f**−**g**) luminescent methods. (**b-e**) THP1-Blue ISG cells were pretreated with (**b**) 50-300 μM NPPB, (**c**) 50-200 μM DIDS, (**d**) 50-200 μM IAA-94, (**e**) 50-200 μM FFA, and 2.5 μM H-151 (STING inhibitor) for 1 h then transfected with 2×10^−2^ μg/µL HT-DNA overnight. Error bars indicate the mean ± standard error of the mean (s.e.m.) of three independent measurements. ^*^*P* < 0.05; ^**^*P* < 0.01; ^****^*P* < 0.0001. One-way analysis of variance (ANOVA) followed by Dunnett’s test for multiple comparison. ns, not significant. (**f-g**) RAW-Dual cells were pretreated with 100 μM NPPB, 150 μM DIDS, and 15 μM H-151 for 1 h then transfected with 2×10^−2^ μg/µL HT-DNA for (**f**) overnight, (**g**)1-24 h. Error bars indicate the mean ± standard error of the mean (s.e.m.) of three independent measurements. ^****^*P* < 0.0001. One-way analysis of variance (ANOVA) followed by Dunnett’s test for multiple comparison. **(h)** IFN-β mRNA expression levels in THP1 cells pretreated with 100 μM NPPB,150 μM DIDS,15μM H-151 for overnight and then transfected with 2×10^−2^ µg/µL HT-DNA overnight. mRNA levels were measured by the RT-qPCR. Error bars indicate the mean ± standard error of the mean (s.e.m.) of three independent measurements. ^***^*P* < 0.001; ^****^*P* < 0.0001. One-way analysis of variance (ANOVA) followed by Dunnett’s test for multiple comparison. **(i)** Western blotting to measure the protein expression level of p-IRF_3_, IRF_3_ and β-actin in THP-1 cells that were pretreated with 100 μM NPPB,150 μM DIDS, and 15 μM H-151 overnight and then transfected with 2×10^−2^ µg/µL HT-DNA overnight. Experiments were performed in three biological replicates. **(j)** Primary human dermal fibroblasts (HDF) were pretreated with 100 μM NPPB, 150 μM DIDS for 18 h and then transfected with 2×10^−2^ µg/µL HT-DNA for 6 h. Cells were fixed using paraformaldehyde and stained with anti-STING antibodies. Experiments were performed in three biological replicates.

The activation of cGAS-STING pathway is accompanied by dramatic IFN-β mRNA induction, which can be monitored by qRT-PCR analysis. To further validate the inhibitory effect of chloride channel blockers on cGAS-STING pathway, we monitored the mRNA levels of IFN-β. The IFN-β mRNA level significantly increased upon HT-DNA stimulation. The enhancement of IFN-β mRNA level was reduced significantly in cells pretreated with NPPB and DIDS (Figure 1h). For further insight into the mechanism of inhibition, we investigated whether chloride homeostasis dysregulation also inhibits the STING to IRF3 signaling. During the immune cascade in response to cytosolic HT-DNA, an important downstream signaling member is IRF3. Through western blot analysis, we observed IRF3 phosphorylation upon HT-DNA stimulation. The IRF3 phosphorylation was reduced in the presence of NPPB and DIDS (Figure 1i, Supplementary Figure 3-4). It was reported that 2’3’-cGAMP binds to STING, triggering STING polymerization and its downstream signaling.^35^ To examine if STING is also inhibited by cellular chloride dysregulation, immunofluorescence imaging was carried out to visualize the cellular distribution of STING in primary human dermal fibroblasts. Upon HT-DNA stimulation, STING-puncta formation occurred, and characteristic large bright puncta were observed. However, samples pretreated with NPPB and DIDS showed collectively less expression of large globular puncta (Figure 1j). These data suggest that chloride homeostasis is essential for the activation of cGAS-STING pathway. Therefore, dysregulation of chloride homeostasis attenuates the cytosolic DNA-stimulated cGAS-STING pathway.

### Chloride homeostasis is crucial to up-stream cGAS-STING signaling

cGAS-STING signaling is quenched by the degradation of activated STING complexes via ubiquitination^85^ and autophagy.^86,87^ Lysosome malfunction reduces the extent of autophagy and causes the enhanced cGAS-STING activation.^88^ Therefore, we sought to find out if the inhibited cGAS-STING signaling is due to the enhanced extent of autophagy. We first validated the inhibitory effect of chloride channel blockers by transfecting the cells with a halide sensitive yellow fluorescent protein (YFP) and incubating the cells with iodide. In this case, the iodide was imported into the cells and able to quench the fluorescence of YFP.^89^ In the presence of chloride channel blockers, iodide transport was inhibited, so the cells displayed a relatively high fluorescence intensity compared to the vehicle (Figure 2a-b). Next, we performed an autophagy analysis to find out the impact of chloride channel blockers on the extent of autophagy. The cells were transfected with RFP-GFP-LC3B tandem sensor, which allows monitoring of autophagy using pH-sensitive green fluorescent protein (GFP) paired with a pH-insensitive red fluorescent protein (RFP).^90^ By measuring fluorescence intensity of GFP (green channel) and RFP (red channel), proper autophagic function can be monitored by anticipating autophagosome-lysosome fusion decreases GFP intensity and RFP intensity. Compared to vehicle control, cells treated with chloroquine (an autophagy/lysosome inhibitor), NPPB, and DIDS, displayed a relatively high fluorescence intensity of RFP and GFP (Figure 2c-e). It suggests that chloroquine, NPPB, and DIDS affect both autophagy and lysosome function. To further evaluate lysosome function, we measured the lysosomal pH upon treatment of chloride channel blockers. We measured the lysosomal pH by labeling the lysosomes with FITC-dextran and TMR-dextran; tetramethylrhodamine (TMR) being a pH insensitive fluorophore functions as the reference while fluorescein isothiocyanate (FITC), a pH sensitive fluorophore, could be used to monitor acidification. Through multichannel imaging, we could measure the lysosomal pH by obtaining the fluorescence intensity ratio (G/R) between FITC (G) and TMR (R) (Figure 2f−g, Supplementary Figure 5). NH_4_Cl treatment functions as the positive control that raises lysosomal pH and gives insight to relative G/R values when lysosomal pH is increased. Upon NH_4_Cl treatment, the G/R ratio of lysosomes increased significantly, which indicates the increase of lysosomal pH. The increased G/R ratio was also observed in the cells undergoing incubation with chloride channel blockers (Figure 2g). This leads us to believe that chloride channel blockers are disrupting lysosomal pH and inhibiting lysosome function.

**Figure 2.**
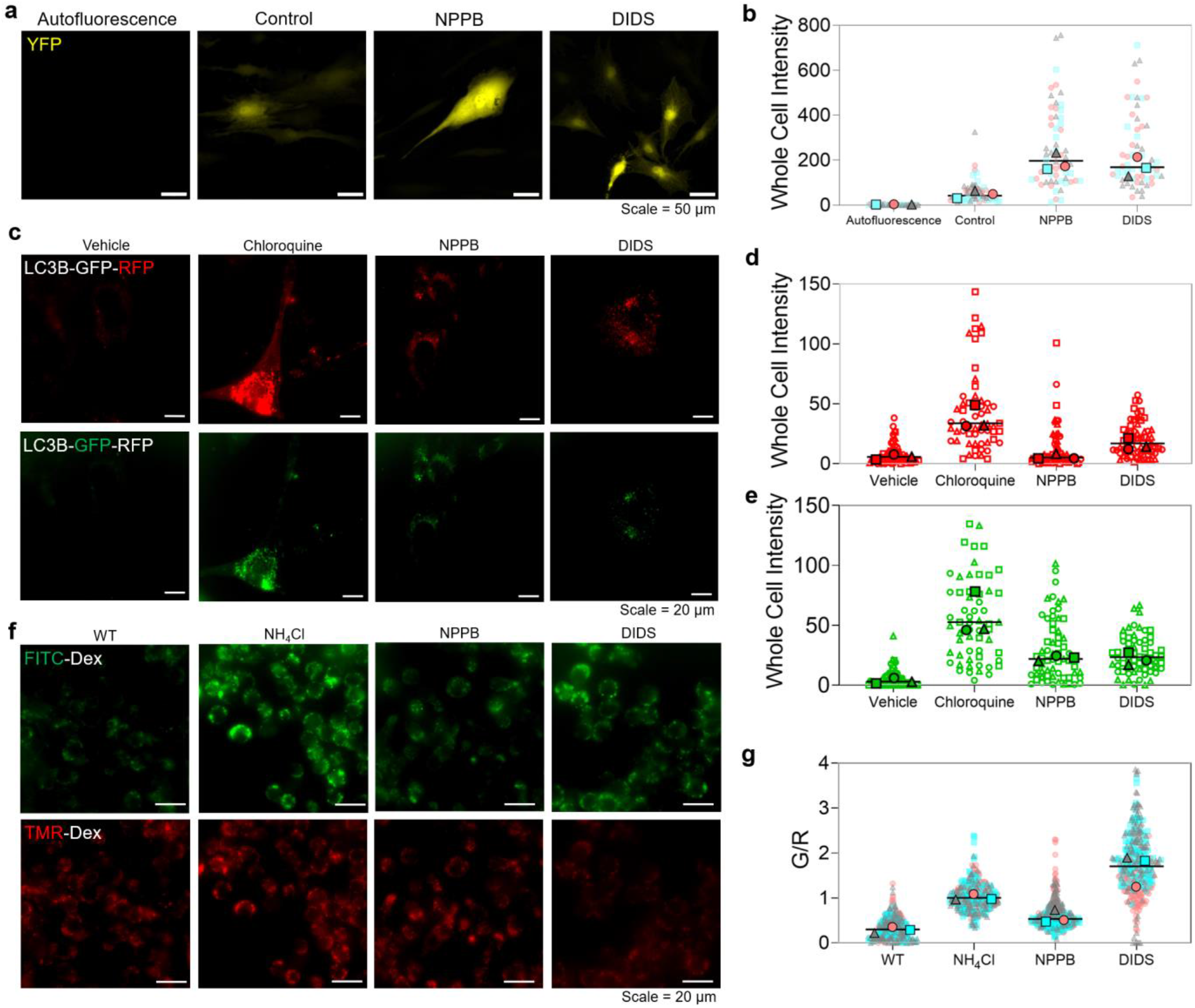
Lysosome function is inhibited by chloride homeostasis dysregulation. **(a)** Representative fluorescence images of primary HDF cells expressing halide sensitive YFP. Cells were incubated with 100 μM NPPB, and 150 μM DIDS overnight before adding the iodide buffer. **(b)** Whole cell fluorescence intensity of YFP-expressing HDF (>20 cells per trial) in the presence or absence of 100 μM NPPB, and 150 μM DIDS. Experiments were performed in triplicate. The median value of each trial is given by a square, circle and triangle symbol. **(c)** Representative fluorescence images of HDF cells transduced with Premo™ tandem autophagy sensor (RFP-GFP-LC3B) upon treatment of 100 μM Chloroquine, 100 μM NPPB, 150 μM DIDS. (**d-e**) Whole cell fluorescence intensity, in (**d**) RFP and (**e**) GFP channels, of HDF cells (>20 cells per trial) transduced with autophagy sensor upon treatment of 100 μM chloroquine, 100 μM NPPB, and 150 μM DIDS. Experiments were performed in triplicate. The median value of each trial is given by a square, circle and triangle symbol. **(f)** Representative fluorescence images of lysosomes of RAW 264.7 macrophage cells labeled with TMR-dextran and FITC dextran in the presence or absence of 1 mM NH_4_Cl, 100 μM NPPB, and 150 μM DIDS. Images acquired in FITC (G) and TMR (R) channels. **(g)** Quantification of lysosomal G/R ratio correlating to (**f**). (>25 cells and >100 lysosomes per trial). Experiments were performed in triplicate. The median value of each trial is given by a square, circle and triangle symbol.

It stands to reason that if autophagy serves as the predominant mechanism to attenuate the cGAS-STING signaling, altering lysosome function would result in inhibited autophagy and enhanced signaling.^91^ However, we didn’t observe the enhanced cGAS-STING signaling in the experiments of Figure 1 even though autophagy was inhibited by chloride channel blockers. Instead, the HT-DNA-induced cGAS-STING signaling was inhibited upon treatment of chloride channel blockers (Figure 1). These results indicate that chloride homeostasis dysregulation attenuates the cGAS-STING signaling in lysosome-independent manner. This leads to the thought that cellular chloride is crucial to up-stream cGAS-STING signaling, prompting the need for thorough investigation.

### Chloride dysregulation attenuates intracellular 2’3’-cGAMP production

To eliminate the possibility that chloride channel blockers reduce the efficiency of HT-DNA transfection and result in inhibited cGAS-STING signaling, we pre-stimulated the cells with HT-DNA and washed the cells with PBS before incubation with chloride channel blockers overnight. In this condition, chloride channel blockers still attenuated the cGAS-STING signaling (Figure 3a). It suggests that neither DIDS nor NPPB inhibits cGAS-STING signaling by blocking the lipofectamine-based HT-DNA transfection. Next, we sought to find out whether chloride homeostasis affects 2’3’-cGAMP production. The enzymatic activity of cGAS is sensitive to ionic conditions.^92^ The impact of various cations on cGAS activity has been extensively studied.^31,92,93^ However, the most prevalent anion in cells, chloride, has been neglected. We employed ELISA to measure 2’3’-cGAMP production in THP-1 cells. Upon HT-DNA stimulation, a significant increase in 2’3’-cGAMP production in THP-1 was observed. The ELISA assay results showed a significant decrease in 2’3’-cGAMP production when the cells were pre-treated with chloride channel blockers NPPB, DIDS, and IAA-94 (Figure 3b). A similar trend was observed for the ELISA assay conducted on RAW 264.7 macrophages (Figure 3c). These results suggest that dysregulation of chloride homeostasis attenuates the HT-DNA stimulated cGAS-STING signaling by suppressing the 2’3’-cGAMP production.

**Figure 3.**
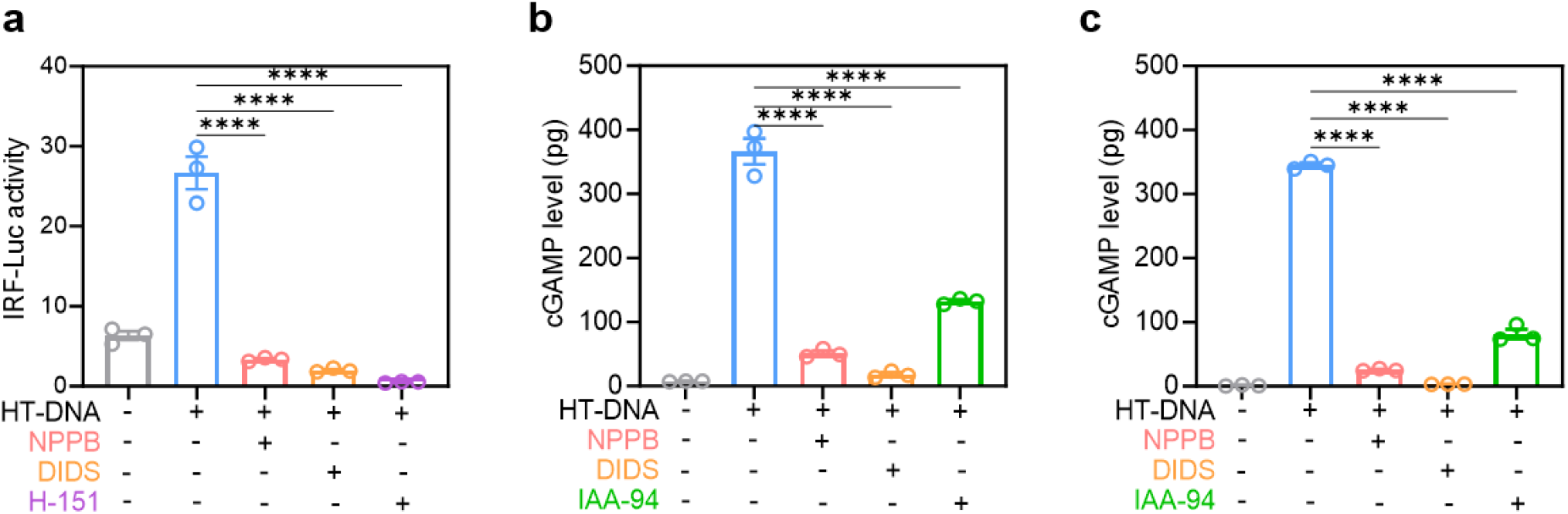
Chloride dysregulation attenuates intracellular 2’3’-cGAMP production. **(a)** RAW-Dual cells were pretreated with 2×10^−2^ μg/µL HT-DNA for 30 min followed by washing using PBS then incubation with 100μM NPPB, 150μM DIDS, 15 μM H-151 overnight. Error bars indicate the mean ± standard error of the mean (s.e.m.) of three independent measurements. ^****^*P* < 0.0001. One-way analysis of variance (ANOVA) followed by Dunnett’s test for multiple comparison. ns, not significant. (**b**-**c**) Intracellular 2’3’-cGAMP level was detected by ELISA. (**b**) THP-1 cells and (**c**) RAW 264.7 cells were pretreated with 100 μM NPPB, 150 μM DIDS, and 200 μM IAA-94 for 24 h then transfected with 2×10^−2^ μg/µL HT-DNA for 6h. Error bars indicate the mean ± standard error of the mean (s.e.m.) of three independent measurements.^****^*P* < 0.0001. One-way analysis of variance (ANOVA) followed by Dunnett’s test for multiple comparison. ns, not significant.

### Dysregulation of chloride homeostasis attenuates STING polymerization and STING-IRF3 signaling

It has been reported that upon activation by 2’3’-cGAMP, STING can function as a proton channel, inducing a pH increase in the Golgi.^71^ As chloride is the most abundant anion in cells and it functions as the counter ion during proton pumping, we anticipated that chloride dysregulation also affects STING. To investigate STING and its downstream signaling, we stimulated the cells with 2’3’-cGAMP to directly activate STING to IRF3 signaling. In this case, the impact of chloride on cGAS could be eliminated. Considering the cellular uptake of 2’3’-cGAMP is through an anion channel, chloride channel blockers may inhibit the STING-IRF3 signaling by suppressing cellular uptake of 2’3’-cGAMP.^94^ Therefore, we stimulated the RAW IRF3 reporter cells with 2’3’-cGAMP, washed the cells and incubated the cells with chloride channel blockers NPPB and DIDS. We observed the STING-IRF3 signaling was significantly reduced upon treatment of NPPB (Figure 4a) and DIDS (Figure 4b). We also employed the murine STING ligand, DMXAA, to activate STING and investigated the STING to IRF3 signaling. We found that DIDS inhibits DMXAA-stimulated STING**-**IRF3 signaling (Figure 4b). We then investigated the STING-IRF3 signaling using THP-1 IRF3 reporter cells. The cells were pre-incubated with chloride channel blockers for 1 h and stimulated with 2’3’-cGAMP overnight (Supplementary Figure 6a−c). Chloride channel blockers inhibited 2’3’-cGAMP-stimulated STING-IRF3 signaling. A comparable trend was observed in the cells upon activation by STING activator MSA-2 (Figure 4c, Supplementary Figure 4d). Chloride channel blockers inhibited MSA-2-stimulated STING-IRF3 signaling. Analysis of IFN-β mRNA levels revealed that both NPPB and DIDS suppressed the interferon response triggered by 2’3’-cGAMP (Figure 4d) and MSA-2 (Figure 4e). Through western blot analysis, we observed the increase of IRF3 phosphorylation when THP-1 cells were stimulated with 2’3’-cGAMP. Upon treatment with NPPB (Figure 4f, Supplementary Figure 7−8) and DIDS (Figure 4g, Supplementary Figure 9−10) in THP-1 cells, IRF3 phosphorylation was notably reduced. These results reveal that chloride homeostasis dysregulation directly attenuates STING-IRF3 signaling.

**Figure 4.**
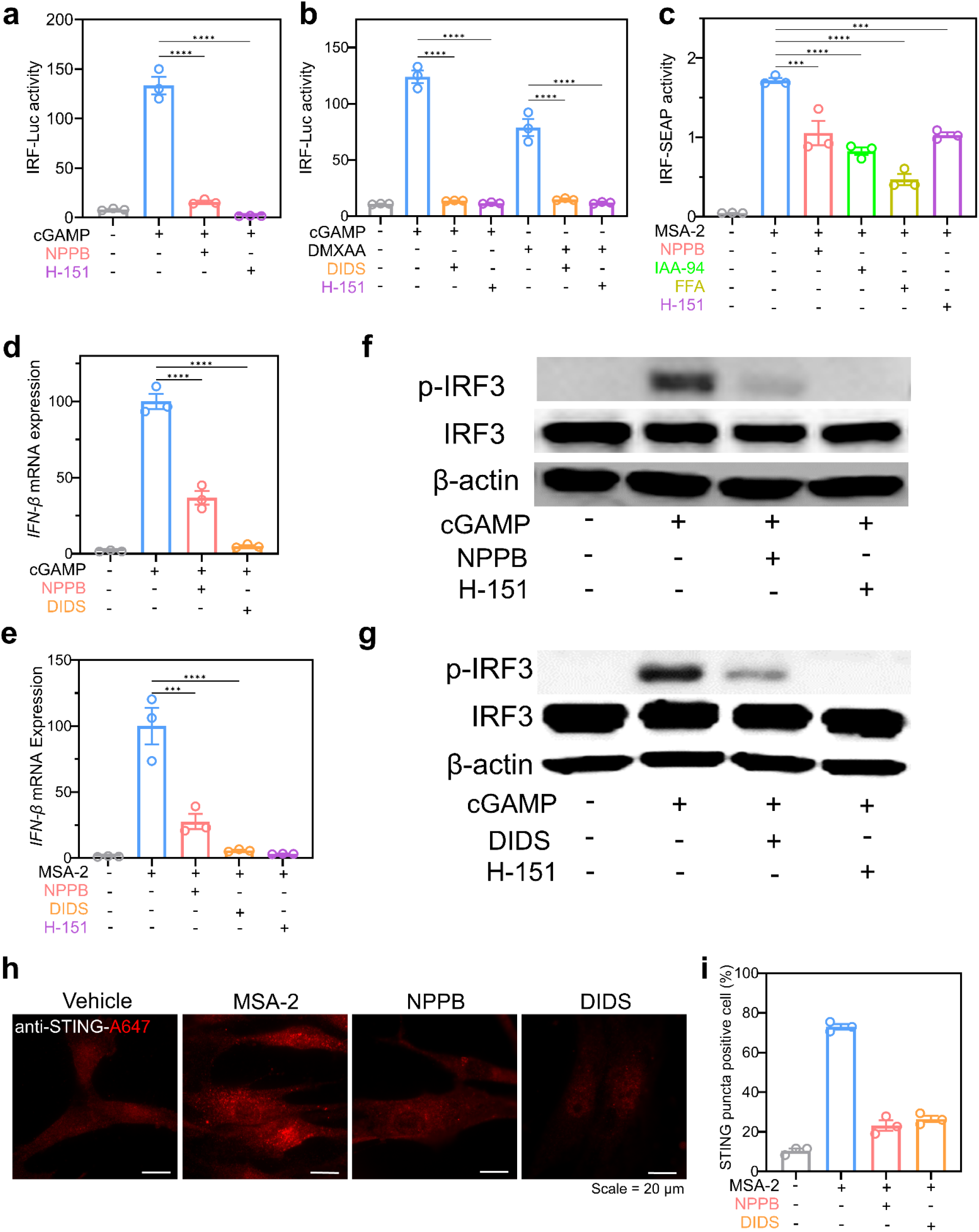
Dysregulation of chloride homeostasis attenuates STING polymerization and STING-IRF_3_ activation. **(a)** RAW-Dual cells were pretreated with 100 µM 2,3-cGAMP for 30 min, followed by washing with PBS, and then incubated with 100 μM NPPB and 15 μM H-151 overnight. **(b)** RAW-Dual cells were pretreated with 100 µM DMXAA or 100 µM 2,3-cGAMP for 30 min, followed by washing with PBS, and then incubated with 150 μM DIDS, and 15 μM H-151 overnight. **(c)** THP1-Blue ISG cells were pretreated with 100 μM NPPB, 200 µM IAA-94, 100 μM FFA, and 15 μM H-151 for 1 h and then stimulated with 30 µM MSA-2 overnight. **(d)** IFN-β mRNA expression levels were measured in THP1 cells pretreated with 100 μM NPPB, and 150 μM DIDS overnight and stimulated with 100 µM 2,3-cGAMP overnight. mRNA levels were measured using the RT-qPCR. **(e)** IFN-β mRNA expression levels were measured in THP1 cells pretreated with 100 μM NPPB, 150 μM DIDS and 15 μM H-151 overnight and stimulated with 30 µM MSA-2 overnight. mRNA levels were measured using the RT-qPCR. **(a-e)** Error bars indicate the mean ± standard error of the mean (s.e.m.) of three independent measurements.^****^*P* < 0.0001. One-way analysis of variance (ANOVA) followed by Dunnett’s test for multiple comparison. ns, not significant. **(f-g)** Western blotting to measure the protein expression level of p-IRF_3_, IRF_3_ and β-actin in THP-1 cells that were pretreated with (**f**) 100 μM NPPB, (**g**) 150 μM DIDS, and 15 μM H-151 overnight and then stimulated with 100 µM 2,3-cGAMP overnight. Experiments were performed in three biological replicates. **(h)** Immunofluorescent analysis of HDF cells pretreated with 100 μM NPPB, and 150 μM DIDS overnight and then stimulated with 30 µM MSA-2 for 6 h. **(i)** Percentage of cells with STING puncta correlating to (**h**). Experiments were performed in three biological replicates (>125 cells per trial). Error bars indicate the mean ± standard error of the mean (s.e.m.) of three independent measurements. ^****^*P* < 0.0001. One-way analysis of variance (ANOVA) followed by Dunnett’s test for multiple comparison.

To further investigate the inhibitory mechanism by chloride homeostasis dysregulation, an immunofluorescence assay was carried out in primary human fibroblast. Human fibroblasts were chosen for this experiment due to their flattened cell morphology, in contrast to the small, round morphology observed in THP-1 cells. The observed augmentation in the quantity of cells harboring STING puncta was notably pronounced following treatment with STING activator MSA-2. Conversely, a marked reduction in this number was observed in cells pre-exposed to chloride channel blockers NPPB and DIDS (Figure 4h−i). All these data collectively suggest that chloride homeostasis dysregulation attenuates STING-IRF3 signaling by inhibiting STING polymerization.

### Non-selective chloride channel blockers suppress the excessive activation of STING signaling in Niemann-Pick disease type C

cGAS-STING pathway is considered as a critical determinant of neuropathophysiology.^95^ To explore the role of chloride homeostasis in cGAS-STING associated pathology, we investigated the impact of non-selective chloride channel blockers on the hyper-activated STING signaling in the Niemann-Pick disease type C (NPC). NPC is a rare genetic disorder that is primarily associated with lysosome dysfunction. There are two subclasses of NPC, NPC type C1 and NPC type C2, each associated with the mutation of either the NPC1 gene or the NPC2 gene, respectively. It was reported that deficiency of NPC1 enhances the STING-IRF3 signaling by blocking the lysosomal degradation.^95^ As we found that dysregulation of chloride homeostasis attenuated the HT-DNA stimulated cGAS-STING signaling, we hypothesized that non-selective chloride channel inhibitors could potentially inhibit the boosted STING signaling in NPC. In order to activate the cGAS-STING pathway, we transfected HT-DNA to primary skin fibroblasts derived from normal individuals and NPC patients. Compared to normal individuals, NPC patient cells transfected with HT-DNA exhibited a markedly elevated mRNA level of IFN−β (Figure 5a, blue bars). This finding aligns with the documented excessive activation of STING signaling observed in NPC1 knockout cells.^95^ We also treated the NPC patient cells with non-selective chloride channel inhibitors NPPB, DIDS, and FFA. Upon treatment of DIDS, the HT-DNA stimulated over-activated STING signaling was inhibited in patient’s cells (Figure 5a, orange bars). However, both NPPB and FFA showed patient-to-patient variability in attenuating HT-DNA stimulated over-activated IFN-β expression (Figure 5a, yellow and red bars). We speculate that the patient-to-patient variation may be due to the difference in NPC1 mutation that causes the different residual protein activity. This result suggests that chloride homeostasis plays an important role in NPC1 deficiency-enhanced STING signaling in NPC. We also investigated the general health of the cells with chloride channel blocker treatment. We found that chloride channel blockers (NPPB, DIDS, IAA-94, FFA) did not cause any cell viability change or induce cell apoptosis (Figure 5b-c and supplementary Figure 2 & 11). All together, these results provide an insight that chloride homeostasis is criticial for the cGAS-STING pathway. By disturbing chloride homeostasis, we are able to suppress the over-activated cGAS-STING signaling in STING associated diseases in which immunomodulation of the type I IFN pathway might show therapeutic benefit.

**Figure 5.**
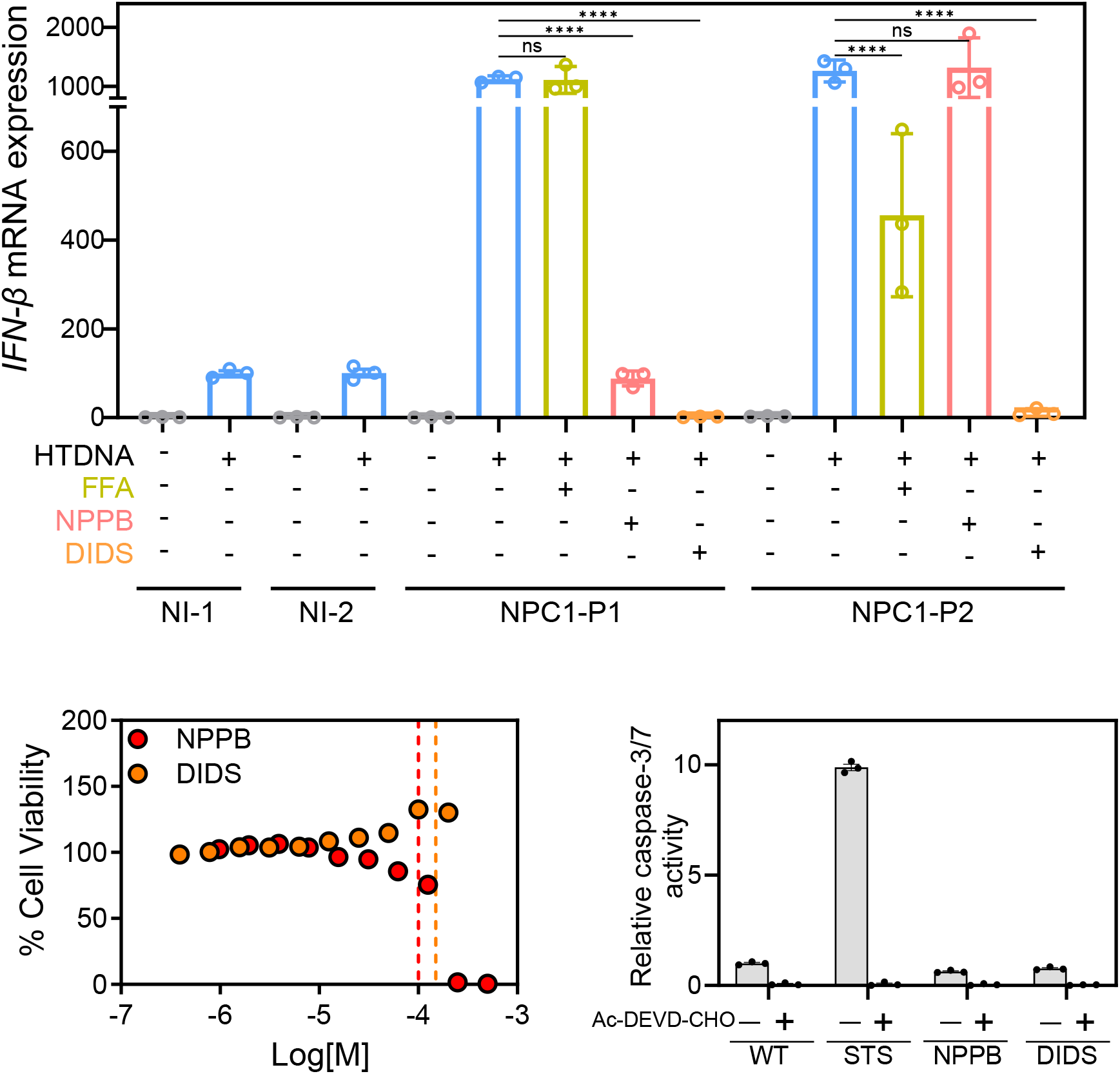
Non-selective chloride channel blockers suppress the excessive activation of STING signaling in Niemann Pick disease type C. **(a)** IFN-β mRNA expression levels were measured in skin fibroblasts from normal individuals (NI) and Niemann Pick Type C1 (NPC1) patients. Cells were pretreated with 100 μM NPPB,150 μM DIDS, and 100 µM FFA overnight and then transfected with 2×10^−2^ µg/µL HT-DNA. mRNA levels were measured using the RT-qPCR. **(b)** Cell viability curve of THP-1 cells upon incubation of NPPB and DIDs overnight. Dashed lines indicate concentrations of compounds used throughout study. **(c)** Caspase-3/7 activity in THP-1 cells. Cells were pretreated with 2 µM staurosporine (STS), 100 µM NPPB, and 150 µM DIDS overnight. 20 µM Ac-DEVD-CHO (caspase inhibitor) was added before analysis as a negative control. (**a**-**c**) Error bars indicate the mean ± standard error of the mean (s.e.m.) of three independent measurements.

## Discussion

The autophagic degradation of activated STING complexes serves as a regulatory brake on the cGAS-STING pathway.^87,88,96^ Therefore, malfunction of lysosomes exacerbates the pathway activation. On the other hand, dysregulation of lysosomal chloride impedes lysosomal function and the activity of lysosomal enzymes.^59,97,98^ Therefore, we initially anticipated that chloride homeostasis would modulate the signaling context through autophagy. Following the treatment of non-selective chloride channel blockers, we noted a reduction in lysosomal function and a corresponding decrease in autophagy (Figure 2c−g). As the lysosomal function was reduced, enhanced cGAS-STING signaling was expected. Contrary to the anticipation that lysosomal chloride dysregulation might enhance the HT-DNA simulated cGAS-STING signaling, we observed a notable attenuation of signaling instead (Figure 1b−j). Therefore, it directed our attention to the possibility that chloride homeostasis dysregulation may inhibit the up-stream cGAS-STING pathway.

Through the 2’3’-cGAMP ELISA assay, we found that chloride dysregulation suppresses the 2’3’-cGAMP production in both THP-1 and RAW 264.7 cells (Figure 3b-c). Meanwhile, by directly activating STING-to-IRF3 signaling using STING activators such as 2’3’-cGAMP and MSA-2, we could investigate the impact of chloride dysregulation on STING-to-IRF3 signaling. The results of IRF3 reporter assay, western blot analysis, and IFNβ mRNA levels analysis (Figure 4a-g) indicated that chloride channel blockers reduced the STING-mediated activation of IRF3.

It has been reported that upon activation and polymerization, STING could function as a proton channel and increase the pH of Golgi.^99^ Chloride channel neutralizes the charge of the proton flux and maintains Golgi membrane potential close to zero.^100^ We anticipated that chloride homeostasis dysregulation may also inhibit this process and cause the inhibition of STING polymerization. Encouragingly, we observed the MSA-2 activated STING polymerization was inhibited by chloride channel blockers (Figure 4h−i). The number of cells containing relatively large STING puncta, induced by STING polymerization, was significantly diminished following chloride channel blocker treatment. It demonstrates that chloride homeostasis dysregulation attenuates STING-to-IRF3 signaling by inhibiting the STING polymerization.

On the other hand, it has been reported that the deficiency of NPC1 enhances the STING-IRF3 signaling by impeding lysosomal degradation. Over-activated STING signaling is imperative for severe neurological degeneration in NPC.^95^ We therefore thought that we could potentially mitigate the NPC pathology by dampening the STING signaling. We initially explored the hyperactivation of the STING pathway in skin fibroblasts derived from patients with NPC. In contrast to cells from normal individuals, the HT-DNA stimulated cGAS-STING signal was significantly elevated in cells obtained from NPC patients (Figure 5a). Our results align with the discovery that NPC1 deficiency results in hyper-activated STING signaling. Finally, we observed that chloride channel blockers displayed different levels of inhibition in cGAS-STING signaling across different NPC patients. The difference in inhibitory effect may be due to the patient-to-patient variation as different NPC1 mutations may lead to varying levels of residual protein activity.

We herein uncover the correlation between chloride homeostasis and cell signaling. We demonstrated that chloride homeostasis is important for cGAS-STING pathway activation. Cellular chloride dysregulation attenuates the HT-DNA stimulated cGAS-STING signaling. We found that chloride homeostasis dysregulation inhibits the 2’3’-cGAMP production and attenuates STING-to-IRF3 signaling by inhibiting the STING polymerization. We also showed the capability of chloride channel blockers to inhibit the over-activated STING signaling in NPC, which contributes to NPC neuropathology. These results provide us an insight into chloride channel-targeted therapy for NPC or diseases that immunomodulation of the type I IFN pathway might show therapeutic benefit.

## Limitations of Study

In this study, we investigated the impact of chloride homeostasis on cGAS-STING signaling. We dysregulated whole cell chloride homeostasis by using the non-selective chloride channel blockers such as NPPB, DIDS, IAA-94, and FFA. We are aware of the likelihood that non-selective chloride channel blockers may exhibit diverse impacts on different chloride channels,^101^ which may be the reason that both 2’3’-cGAMP production and STING-to-IRF3 signaling are attenuated by the non-selective chloride channel blockers. Functional investigation of chloride channels will be conducted through genetic manipulation in the subsequent projects. It will find out which chloride channel regulates 2’3’-cGAMP production or STING-to-IRF3 signaling and identify the inhibition mechanism.

We are aware that YFP is a pH-sensitive, non-quantitative tool to monitor the intracellular chloride and another pH-independent lysosomal chloride reporter was reported.^102^ However, the Premo™ YFP halide sensor is the only commercially available tool for chloride imaging. Therefore, we used it to validate the chloride homeostasis dysregulation induced by the non-selective chloride channel blockers.

## Methods

### Chemicals

NPPB, DIDS, H-151, IAA-94, FFA, MSA-2, DMXAA, staurosporine, and Ac-DEVD-CHO were purchased from Cayman Chemicals (Ann Arbor, MI, USA). 2’3’-cGAMP were purchased from Chemietek (Indianapolis, IN, USA). Deoxyribonucleic acid sodium salt from herring testes (HT-DNA) was purchased from Sigma-Aldrich (St Louis, MA, USA). Lipofectamine 3000 transfection kit was purchased from InvivoGen (San Diego, CA, USA). Phospho-IRF3(Ser396) [4D4G], β-actin [8H10D10] primary antibodies and anti-rabbit-HRP or anti-mouse-HRP antibody were purchased from Cell Signaling Technology (Danvers, MA, USA). IRF3 [A19717] were purchased from ABclonal (Woburn, MA, USA).

### Cell lines and culturing

THP1-Blue™ ISG cells were purchased from InvivoGen (San Diego, CA, USA). THP1 cells were purchased from ATCC (Manassas, VA, USA). These cell lines were routinely maintained in RPMI 1640 containing 10% heat inactivated FBS, 10 mM HEPES,1mM sodium pyruvate, 4500 mg/L glucose, and 1500 mg/L sodium bicarbonate, 100 U/mL penicillin, 100 mg/mL streptomycin. RAW-Dual™ cells were purchased from InvivoGen (San Diego, CA, USA). RAW 264.7 cells were purchased from ATCC (Manassas, VA, USA). These cell lines were routinely maintained in Dulbecco’s Modified Eagle’s Medium (DMEM) supplemented with 10% heat-inactivated fetal bovine serum, Pen-Strep (100 U/mL-100 µg/mL). THP1-Blue™ ISG cells and RAW-Dual™ cell lines were selectively pressured using growth media containing Zeocin® according to the manufacturer’s protocol. Primary Human Dermal Fibroblasts (HDF) were purchased from ATCC (Manassas, VA, USA). This cell line was routinely maintained in Dulbecco’s Modified Eagle’s Medium/ Nutrient mixture F-12 (DMEM/F12) supplemented with 10% fetal bovine serum, Pen-Strep (100 U/mL-100 µg/mL). DMEM/F12 was purchased from (Thermo Fisher Scientific, CA, USA). DMEM, RPMI 1640 and fetal bovine serum was purchased from Corning (Corning, NY, USA). Cell lines were cultured in 37 °C with 5% CO2 atmosphere. Fibroblasts GM17913, GM18388 and GM22277 were purchased from Coriell Institute (Camden, NJ) and cultured with the suggested protocols from Coriell Institute.

### Western blot

Chloride inhibitors effect on p-IRF3 expression was tested on human leukemia monocytic THP-1 cells. 1×10^6^ THP-1 cells were seeded in a 24-well dish. 1 h later, cells were treated 100 µM NPPB, 150 µM DIDS overnight. Followed by 100 µM 2’3’-cGAMP or transfected with 2x10^−2^ µg/µL HT-DNA via Lipofectamine 3000 reagent. After the indicated time periods, cells were collected by centrifugation and total protein extracts were obtained using M-PER™ Mammalian Protein Extraction Reagent (Thermo-Fisher Scientific, CA, USA) supplemented with pierce protease kit and phosphatase inhibitors (Thermo Fisher Scientific, CA, USA). The cell lysates were centrifuged at 14,000 × g for 10 min at 4°C and the protein containing supernatant was collected. Protein concentration of each sample was determined by the Pierce™ Rapid Gold BCA Protein Assay Kit (Thermo Fisher Scientific, CA, USA). Equal amounts of proteins (up to 20 μg) were separated on SDS-PAGE gel and transferred to a nitrocellulose membrane. The membrane was then blocked with 1× TBST (Tris-buffered saline, 0.1% Tween 20 (20 mM Tris pH7.5, 150 mM NaCl and 0.1% Tween 20)) for 1 h at room temperature. After blocking, the membrane was incubated with primary antibodies (Phospho-IRF3(Ser396), IRF3 and β-actin) overnight at 4°C. The primary antibody IRF3 was purchased from ABclonal (Woburn, MA, USA). The primary antibodies p-IRF3 and β-actin were purchased from Cell Signaling (Danvers, MA, USA). The membrane was washed by 1× TBST 3 times and incubated with a secondary anti-rabbit-HRP or anti-mouse antibody-HRP (Cell Signaling, Danvers, MA) for 1 h at room temperature. The membrane was washed again, and the signal was detected by SuperSignal™ West Pico PLUS Chemiluminescent Substrate (Thermo-Fisher Scientific, CA, USA).

### Quanti-Blue™ reporter assay

2×10^5^ THP1-Blue™ were seeded in well plate. 1 h later, cells were treated with 100 µM NPPB, 150 µM DIDS, 200 µM IAA-94, 100 µM FFA and 15 µM H-151 for 1 h. Followed by transfection of 2×10^−2^ µg/µL HT-DNA via Lipofectamine 3000 reagent or 100 µM 2’3’-cGAMP or 30 µM MSA-2 overnight. Each condition was replicated in triplicate. The Quanti-Blue™ colorimetric assay InvivoGen (San Diego, CA, USA) was performed according to the manufacturers’ protocol. Plates were read using SpectraMax™ i3/i3x multi-mode plate reader.

### Quanti-Luc™ reporter assay

2×10^5^ RAW-Dual macrophage were seeded in well plate. 24 h later, cells were treated with 100 µM NPPB, 150 µM DIDS, 200 µM IAA-94, and 15 µM H-151 for 1 h. Followed by transfection of 2x10^−2^ µg/µL HT-DNA via Lipofectamine 3000 reagent or 100 µM 2’3’-cGAMP or 100 µM DMXAA overnight. Each condition was replicated in triplicate. The Quanti-LUC™ luminescent assay InvivoGen (San Diego, CA, USA) was performed according to the manufacturers’ protocol. Plates were read using SpectraMax™ i3/i3x multi-mode plate reader.

### ELISA

The effect of chloride inhibitors on 2,3-cGAMP production was tested on human leukemia monocytic THP-1 cells. 1 x 10^6^ THP-1 or RAW 264.7 cells were seeded in well plate. 1h later, cells were treated 100 µM NPPB, 150 µM DIDS, 200 µM IAA-94 overnight. Followed by transfection with 2x10^−2^ µg/µL HT-DNA via Lipofectamine 3000 reagent InvivoGen (San Diego, CA, USA) for 6 h. Cell pellets were lysed in M-PER™ Mammalian Protein Extraction Reagent (Thermo-Fisher Scientific, CA, USA) supplemented with cOmplete protease inhibitor cocktail (Roche). Lysed cells were centrifuged for 10 min at 14,000 g and 4 °C. Protein concentration of each lysate was measured using the Pierce™ Rapid Gold BCA Protein Assay Kit (Thermo Fisher Scientific, CA, USA). The 2′-3′-cGAMP content were measured by 2′-3′-cGAMP ELISA kit, Cayman Chemicals (Ann Arbor, MI, USA) according to the manufacturer’s instructions. Plates were read using SpectraMax™ i3/i3x Multi-Mode plate reader.

### Wide field microscopy

All wide field imaging was done using an IX-83 inverted microscope (Olympus Corporation of the Americas, Center Valley, PA, USA) and Prime BSI CMOS camera (Photometrics, USA). Filter cubes, shutter and CMOS camera were controlled using Cellsens Dimension 4.1 (Olympus Corporation of the Americas, Center Valley, PA, USA).

### Immunofluorescence

Primary Human Dermal Fibroblasts (HDF) were plated on 35mm Glass Bottomed Dishes (CellVis, Mountain view, CA, USA). Cells were treated with 100 µM NPPB, 150 µM DIDS overnight. Followed by stimulation using 30 µM MSA-2 or transfected with 2×10^−2^ µg/µL HT-DNA via Lipofectamine 3000 reagent. Cells were then fixed using 4% paraformaldehyde (Thermo Fisher Scientific, CA, USA), permeabilized using 0.1% Triton-X-100 (Thermo Fisher Scientific, CA, USA), followed by blocking using 2% BSA in PBS. They were then incubated overnight at 4 °C with the primary antibody (anti-TMEM173/STING antibody (OT14H1),1:100 dilution) (Invitrogen, MA,USA). After they were rinsed with PBS-T, then were incubated with the secondary antibody (Alexa-Fluor®-647-conjugated goat anti-mouse IgG H+L,1:2000 dilution) for 60 min at room temperature and counterstained with DAPI (Sigma-Aldrich) for another 10 min. The cells were washed twice with PBS-T and observed using a 100×, 1.45 NA, apochromat oil immersion objective (UPLANXAPO, Olympus Corporation of the Americas, Center Valley, PA, USA). Alexa 647 channel images were obtained using a filter cube containing 620/60 band pass excitation filter, 700/75 band pass emission filter and T660lpxr dichroic filter (Chroma, Bellows Falls, VT, USA). Background subtraction was done for all cells by taking mean intensity over an adjacent cell free area using ImageJ (NIH). STING positive puncta were determined by setting each image to equal contrast levels (in respect to positive control) and visually examining for the presence of large, condensed puncta, in each condition. All Images were taken the same day and were acquired under the same acquisition settings.

### Autophagy Analysis

Primary Human Dermal Fibroblasts (HDF) were plated on 35mm Glass Bottomed Dishes (Cell Vis, Mountain view, CA, USA) and treated with 100 µM NPPB, 150 µM DIDS overnight. Following treatment cells were then transduced with Premo™ Autophagy tandem sensor Red Fluorescent Protein-Green Fluorescent Protein (RFP-GFP)-LC3B (Life Technologies, Invitrogen) and chloroquine was used as control, according to manufacturer’s protocol. Cells were observed using a 100×, 1.45 NA, apochromat oil immersion objective (UPLANXAPO, Olympus Corporation of the Americas, Center Valley, PA, USA). GFP channel images (G) were obtained using a filter cube containing 480/30 band pass excitation filter, 535/40 band pass emission filter and a 505dc dichroic filter. RFP channel images (R) were obtained using a filter cube containing 545/25 band pass excitation filter, 605/70 band pass emission filter and a BS565 dichroic filter (Chroma, Bellows Falls, VT, USA). Background subtraction was done for all images by taking mean intensity over an adjacent cell free area and whole cell intensity was calculated for each cell by taking mean intensity of defined cell specific ROIs on designated channels and calculated using ImageJ (NIH). All Images were taken the same day and were acquired under the same acquisition settings.

### Whole Cell Chloride Analysis

Primary Human Dermal Fibroblasts (HDF) were plated on 35mm Glass Bottomed Dishes (Cell Vis, Mountain view, CA,USA). Cells were transduced with Premo™ Halide Sensor (Life Technologies, Invitrogen) according to manufacturer’s protocol and then incubated with 100 µM NPPB, 150 µM DIDS overnight. Cells were observed using a 60×, 1.42 NA, apochromat oil immersion objective (UPLANXAPO, Olympus Corporation of the Americas, Center Valley, PA, USA). YFP channel images were obtained using a filter cube containing 490/40 band pass excitation filter, ET535/30 band pass emission filter and T515LPXR dichroic filter (Chroma, Bellows Falls, VT, USA). Background subtraction was done for all images by taking mean intensity over an adjacent cell free area and whole cell intensity was calculated for each cell by taking mean intensity of defined cell specific ROIs and calculated using ImageJ (NIH). All Images were taken the same day and were acquired under the same acquisition settings.

### Lysosomal pH analysis

RAW 264.7 murine macrophages were plated on 35mm Glass Bottomed Dishes (Cell Vis, Mountain view, CA, USA) and treated with 1 mM NH_4_Cl (positive control), 100 µM NPPB, 150 µM DIDS overnight. Following treatment cells were then pulsed for 1 h with 1 mg/mL FITC-Dextran (10k MW)/ TMR-Dextran (10k MW) (Sigma-Aldrich, St Louis, MA, USA) diluted in OPTI-MEM and then washed with PBS followed by addition of growth media. Cells were then chased (incubated 37°C with 5% CO2 atmosphere) for 6 h before imaging. Cells were observed using a 100×, 1.45 NA, apochromat oil immersion objective (UPLANXAPO, Olympus Corporation of the Americas, Center Valley, PA, USA). FITC channel images (G) were obtained using a filter cube containing 480/30 band pass excitation filter, 535/40 band pass emission filter and a 505dc dichroic filter. TMR channel images (R) were obtained using a filter cube containing 545/25 band pass excitation filter, 605/70 band pass emission filter and a BS565 dichroic filter (Chroma, Bellows Falls, VT, USA). Background subtraction was done for all images by taking mean intensity over an adjacent cell free area. Lysosome based intensities in (G) and (R) channels were calculated by defining specific lysosome-based ROIs and measuring mean intensity using ImageJ (NIH). Each ROI was carefully defined in the pH insensitive (R) channel. R and G values measured were then subsequently used for ratio metric (R/G) based analysis to interpret relative pH change. All Images were taken the same day and were acquired under the same acquisition settings.

### RT-qPCR

The effect of chloride inhibitors on IFN-β production was tested on human leukemia monocytic THP-1 cells. 1×10^6^ THP-1 were seeded in a 12-Well dish. 1 h later, cells were treated 100 µM NPPB, 150 µM DIDS, 15 µM H-151 overnight. Followed by transfection with 2×10^−2^ µg/µL HT-DNA via Lipofectamine 3000 reagent InvivoGen (San Diego, CA, USA) or 100 µM 2’3’-cGAMP for 18 h. Post stimulation, cells were harvested, and total RNA was extracted by Aurum Total RNA Mini Kit (Bio-Rad,Hercules, California, USA).). Using SuperScript II reverse transcriptase random hexamer primer total RNA was reverse transcribed into cDNA. Real-time PCR was carried out via a QuantiTect SYBR Green PCR Kits and ran on an Axygen MaxyGene II Thermal Cycler. The human IFN-β primer used is listed in Supplementary Table 2. IFN-β Cq values and GAPDH Cq values were used to calculate ΔCq (Cq(IFN-β) - Cq(ref)). Relative gene expression (IFN-β) was obtained using a ΔΔCq calculation method. This is done by normalization of a specific gene target (IFN-β) with experimental treatment to a reference gene (GAPDH).

The effect of chloride inhibitors on IFN-β production was tested on primary patient skin fibroblasts. 2 x 10^6^ were seeded in a 6-Well dish. 18 h later, cells were treated 100 µM NPPB, 150 µM DIDS, 100 µM FFA overnight. Followed by transfection with 2×10^−2^ µg/µL HT-DNA via Lipofectamine 3000 reagent InvivoGen (San Diego, CA, USA) for 18 h. DMSO was used as a negative control. Post stimulation, cells were harvested, and total RNA was extracted by Aurum Total RNA Mini Kit (Bio-Rad,Hercules, California, USA).Using SuperScript II reverse transcriptase random hexamer primer total RNA was reverse transcribed into cDNA. Real-time PCR was carried out via a QuantiTect SYBR Green PCR Kits and ran on an Axygen MaxyGene II Thermal Cycler. The human IFN-β primer used is listed in Supplementary Table 2. IFN-β Cq values and GAPDH Cq values were used to calculate ΔCq (Cq(IFN-β) - Cq(ref)). Relative gene expression (IFN-β) was obtained using a ΔΔCq calculation method. This is done by normalization of a specific gene target (IFN-β) with experimental treatment to a reference gene (GAPDH).

### Cell viability assay

0.1−0.4×10^5^ cells were seeded in 96 well culture plate overnight. Cells were treated with NPPB, DIDS, IAA-94, FFA for 18 h. The CellTiter-Blue® Cell Viability Assay was then performed according to the manufacturers’ protocol (Promega, Madison, Wi, USA). Plates were read using SpectraMax™ i3/i3x multi-mode plate reader.

### Caspase 3/7 activity

0.4×10^5^ cells were seeded in 96 well culture plate overnight. Cells were treated with 100 µM NPPB, 150 µM DIDS, 2 µM Staurosporine (STS) for 18 h. Caspase-Glo® 3/7 Assay was then performed according to the manufacturers’ protocol (Promega, Madison, Wi, USA) and 20 µM Ac-DEVD-CHO was added to select wells for each condition as a negative control. Plates were read using SpectraMax™ i3/i3x multi-mode plate reader.

## Acknowledgements

This work was supported by NIH grants R35GM147112 (K.L.) & R35GM147112-02S2 (K.L.) and Clarkson University start-up fund.

## Author contributions

J.M., H.S., M.W., and K.L. designed the project. J.M., D.W., S.M., D.W., and C. H. performed experiments. J.M., D.W., S.M., D.W., C.C.D., H.S., M.W., and K.L. analyzed the data. J.M., H.S., M.W. and K.L. wrote the paper. All authors discussed the results and gave inputs on the manuscript.

## References

1. Hopfner, K.-P. & Hornung, V. Molecular mechanisms and cellular functions of cGAS-STING signalling. Nat. Rev. Mol. Cell Biol. 21, 501–521 (2020).

2. Stetson, D. B. & Medzhitov, R. Recognition of cytosolic DNA activates an IRF3-dependent innate immune response. Immunity 24, 93–103 (2006).

3. Sun, L., Wu, J., Du, F., Chen, X. & Chen, Z. J. Cyclic GMP-AMP synthase is a cytosolic DNA sensor that activates the type I interferon pathway. Science 339, 786–791 (2013).

4. Banani, S. F., Lee, H. O., Hyman, A. A. & Rosen, M. K. Biomolecular condensates: organizers of cellular biochemistry. Nat. Rev. Mol. Cell Biol. 18, 285–298 (2017).

5. Kranzusch, P. J. et al. Ancient Origin of cGAS-STING Reveals Mechanism of Universal 2’,3’ cGAMP Signaling. Mol. Cell 59, 891–903 (2015).

6. Corrales, L. et al. Direct activation of STING in the tumor microenvironment leads to potent and systemic tumor regression and immunity. Cell Rep. 11, 1018–1030 (2015).

7. Samson, N. & Ablasser, A. The cGAS-STING pathway and cancer. Nat. Cancer 3, 1452–1463 (2022).

8. Domizio, J. D. et al. The cGAS-STING pathway drives type I IFN immunopathology in COVID-19. Nature 603, 145–151 (2022).

9. Gao, D. et al. Activation of cyclic GMP-AMP synthase by self-DNA causes autoimmune diseases. Proc Natl Acad Sci USA 112, E5699–705 (2015).

10. Gray, E. E., Treuting, P. M., Woodward, J. J. & Stetson, D. B. Cutting Edge: cGAS Is Required for Lethal Autoimmune Disease in the Trex1-Deficient Mouse Model of Aicardi-Goutières Syndrome. J. Immunol. 195, 1939–1943 (2015).

11. Gall, A. et al. Autoimmunity initiates in nonhematopoietic cells and progresses via lymphocytes in an interferon-dependent autoimmune disease. Immunity 36, 120–131 (2012).

12. Ablasser, A. et al. TREX1 deficiency triggers cell-autonomous immunity in a cGAS-dependent manner. J. Immunol. 192, 5993–5997 (2014).

13. Decout, A., Katz, J. D., Venkatraman, S. & Ablasser, A. The cGAS-STING pathway as a therapeutic target in inflammatory diseases. Nat. Rev. Immunol. 21, 548–569 (2021).

14. Gulen, M. F. et al. cGAS-STING drives ageing-related inflammation and neurodegeneration. Nature 620, 374–380 (2023).

15. Xie, X. et al. Activation of innate immune cGAS-STING pathway contributes to Alzheimer’s pathogenesis in 5×FAD mice. Nat. Aging 3, 202–212 (2023).

16. Microglial cGAS-STING links innate immunity and Alzheimer’s disease. Nat. Aging 3, 155–156 (2023).

17. Chitnis, T. & Weiner, H. L. CNS inflammation and neurodegeneration. The Journal of Clinical Investigation (2017).

18. Ablasser, A. et al. cGAS produces a 2’-5’-linked cyclic dinucleotide second messenger that activates STING. Nature 498, 380–384 (2013).

19. Gao, P. et al. Cyclic [G(2’,5’)pA(3’,5’)p] is the metazoan second messenger produced by DNA-activated cyclic GMP-AMP synthase. Cell 153, 1094–1107 (2013).

20. Diner, E. J. et al. The innate immune DNA sensor cGAS produces a noncanonical cyclic dinucleotide that activates human STING. Cell Rep. 3, 1355–1361 (2013).

21. Kranzusch, P. J. cGAS and CD-NTase enzymes: structure, mechanism, and evolution. Curr. Opin. Struct. Biol. 59, 178–187 (2019).

22. Kranzusch, P. J., Lee, A. S.-Y., Berger, J. M. & Doudna, J. A. Structure of human cGAS reveals a conserved family of second-messenger enzymes in innate immunity. Cell Rep. 3, 1362–1368 (2013).

23. Civril, F. et al. Structural mechanism of cytosolic DNA sensing by cGAS. Nature 498, 332–337 (2013).

24. Li, X. et al. Cyclic GMP-AMP synthase is activated by double-stranded DNA-induced oligomerization. Immunity 39, 1019–1031 (2013).

25. Zhang, X. et al. The cytosolic DNA sensor cGAS forms an oligomeric complex with DNA and undergoes switch-like conformational changes in the activation loop. Cell Rep. 6, 421–430 (2014).

26. Andreeva, L. et al. cGAS senses long and HMGB/TFAM-bound U-turn DNA by forming protein-DNA ladders. Nature 549, 394–398 (2017).

27. Gentili, M. et al. The N-Terminal Domain of cGAS Determines Preferential Association with Centromeric DNA and Innate Immune Activation in the Nucleus. Cell Rep. 26, 2377-2393.e13 (2019).

28. Luecke, S. et al. cGAS is activated by DNA in a length-dependent manner. EMBO Rep. 18, 1707–1715 (2017).

29. Herzner, A.-M. et al. Sequence-specific activation of the DNA sensor cGAS by Y-form DNA structures as found in primary HIV-1 cDNA. Nat. Immunol. 16, 1025–1033 (2015).

30. Mankan, A. K. et al. Cytosolic RNA:DNA hybrids activate the cGAS-STING axis. EMBO J. 33, 2937–2946 (2014).

31. Du, M. & Chen, Z. J. DNA-induced liquid phase condensation of cGAS activates innate immune signaling. Science 361, 704–709 (2018).

32. Xie, W. et al. Human cGAS catalytic domain has an additional DNA-binding interface that enhances enzymatic activity and liquid-phase condensation. Proc Natl Acad Sci USA 116, 11946–11955 (2019).

33. Wu, J. et al. Cyclic GMP-AMP is an endogenous second messenger in innate immune signaling by cytosolic DNA. Science 339, 826–830 (2013).

34. Zhang, X. et al. Cyclic GMP-AMP containing mixed phosphodiester linkages is an endogenous high-affinity ligand for STING. Mol. Cell 51, 226–235 (2013).

35. Ergun, S. L., Fernandez, D., Weiss, T. M. & Li, L. STING polymer structure reveals mechanisms for activation, hyperactivation, and inhibition. Cell 178, 290-301.e10 (2019).

36. Burdette, D. L. et al. STING is a direct innate immune sensor of cyclic di-GMP. Nature 478, 515–518 (2011).

37. Ishikawa, H. & Barber, G. N. STING is an endoplasmic reticulum adaptor that facilitates innate immune signalling. Nature 455, 674–678 (2008).

38. Sun, W. et al. ERIS, an endoplasmic reticulum IFN stimulator, activates innate immune signaling through dimerization. Proc Natl Acad Sci USA 106, 8653–8658 (2009).

39. Ablasser, A. Structures of STING protein illuminate this key regulator of inflammation. Nature 567, 321–322 (2019).

40. Zhong, B. et al. The adaptor protein MITA links virus-sensing receptors to IRF3 transcription factor activation. Immunity 29, 538–550 (2008).

41. Ishikawa, H., Ma, Z. & Barber, G. N. STING regulates intracellular DNA-mediated, type I interferon-dependent innate immunity. Nature 461, 788–792 (2009).

42. Abe, T. et al. STING recognition of cytoplasmic DNA instigates cellular defense. Mol. Cell 50, 5–15 (2013).

43. Li, X.-D. et al. Pivotal roles of cGAS-cGAMP signaling in antiviral defense and immune adjuvant effects. Science 341, 1390–1394 (2013).

44. Gao, D. et al. Cyclic GMP-AMP synthase is an innate immune sensor of HIV and other retroviruses. Science 341, 903–906 (2013).

45. Kuchitsu, Y. et al. STING signalling is terminated through ESCRT-dependent microautophagy of vesicles originating from recycling endosomes. Nat. Cell Biol. 25, 453–466 (2023).

46. Bai, J. & Liu, F. The cGAS-cGAMP-STING Pathway: A Molecular Link Between Immunity and Metabolism. Diabetes 68, 1099–1108 (2019).

47. Qiao, J. T. et al. Activation of the STING-IRF3 pathway promotes hepatocyte inflammation, apoptosis and induces metabolic disorders in nonalcoholic fatty liver disease. Metab. Clin. Exp. 81, 13–24 (2018).

48. Bao, T., Liu, J., Leng, J. & Cai, L. The cGAS-STING pathway: more than fighting against viruses and cancer. Cell Biosci. 11, 209 (2021).

49. Sze, A. et al. Host restriction factor SAMHD1 limits human T cell leukemia virus type 1 infection of monocytes via STING-mediated apoptosis. Cell Host Microbe 14, 422–434 (2013).

50. Benmerzoug, S. et al. STING-dependent sensing of self-DNA drives silica-induced lung inflammation. Nat. Commun. 9, 5226 (2018).

51. Dou, Z. et al. Cytoplasmic chromatin triggers inflammation in senescence and cancer. Nature 550, 402–406 (2017).

52. Glück, S. et al. Innate immune sensing of cytosolic chromatin fragments through cGAS promotes senescence. Nat. Cell Biol. 19, 1061–1070 (2017).

53. Dmitrieva, N. I. & Burg, M. B. High NaCl promotes cellular senescence. Cell Cycle 6, 3108–3113 (2007).

54. Lee, H. J. et al. Chloride channel accessory 1 integrates chloride channel activity and mTORC1 in aging-related kidney injury. Aging Cell 20, e13407 (2021).

55. Suh, K. S. et al. The organellular chloride channel protein CLIC4/mtCLIC translocates to the nucleus in response to cellular stress and accelerates apoptosis. J. Biol. Chem. 279, 4632–4641 (2004).

56. Ko, S.-K. et al. Synthetic ion transporters can induce apoptosis by facilitating chloride anion transport into cells. Nat. Chem. 6, 885–892 (2014).

57. Miura, G. Death by ions. Nat. Chem. Biol. 10, 795–795 (2014).

58. Lim, C.-Y. & Zoncu, R. The lysosome as a command-and-control center for cellular metabolism. J. Cell Biol. 214, 653–664 (2016).

59. Chakraborty, K., Leung, K. & Krishnan, Y. High lumenal chloride in the lysosome is critical for lysosome function. eLife 6, e28862 (2017).

60. Maliougina, M. & El Hiani, Y. TRPM2: bridging calcium and ROS signaling pathways-implications for human diseases. Front. Physiol. 14, 1217828 (2023).

61. Sterea, A. M., Almasi, S. & El Hiani, Y. The hidden potential of lysosomal ion channels: A new era of oncogenes. Cell Calcium 72, 91–103 (2018).

62. Lytle, C. & Forbush, B. Regulatory phosphorylation of the secretory Na-K-Cl cotransporter: modulation by cytoplasmic Cl. Am. J. Physiol. 270, C437–48 (1996).

63. Curran, M. J. & Brodwick, M. S. Ionic control of the size of the vesicle matrix of beige mouse mast cells. J. Gen. Physiol. 98, 771–790 (1991).

64. Kaila, K., Price, T. J., Payne, J. A., Puskarjov, M. & Voipio, J. Cation-chloride cotransporters in neuronal development, plasticity and disease. Nat. Rev. Neurosci. 15, 637–654 (2014).

65. Seja, P. et al. Raising cytosolic Cl-in cerebellar granule cells affects their excitability and vestibulo-ocular learning. EMBO J. 31, 1217–1230 (2012).

66. Leung, K., Chakraborty, K., Saminathan, A. & Krishnan, Y. A DNA nanomachine chemically resolves lysosomes in live cells. Nat. Nanotechnol. 14, 176–183 (2019).

67. Graves, A. R., Curran, P. K., Smith, C. L. & Mindell, J. A. The Cl-/H+ antiporter ClC-7 is the primary chloride permeation pathway in lysosomes. Nature 453, 788–792 (2008).

68. Kasper, D. et al. Loss of the chloride channel ClC-7 leads to lysosomal storage disease and neurodegeneration. EMBO J. 24, 1079–1091 (2005).

69. Jentsch, T. J., Stein, V., Weinreich, F. & Zdebik, A. A. Molecular structure and physiological function of chloride channels. Physiol. Rev. 82, 503–568 (2002).

70. Bonam, S. R., Wang, F. & Muller, S. Lysosomes as a therapeutic target. Nat. Rev. Drug Discov. 18, 923–948 (2019).

71. Liu, B. et al. Human STING is a proton channel. Science 381, 508–514 (2023).

72. Aichbichler, B. W., Zerr, C. H., Santa Ana, C. A., Porter, J. L. & Fordtran, J. S. Proton-pump inhibition of gastric chloride secretion in congenital chloridorrhea. N. Engl. J. Med. 336, 106–109 (1997).

73. Osei-Owusu, J. et al. Molecular determinants of pH sensing in the proton-activated chloride channel. Proc Natl Acad Sci USA 119, e2200727119 (2022).

74. Yang, J. et al. PAC, an evolutionarily conserved membrane protein, is a proton-activated chloride channel. Science 364, 395–399 (2019).

75. Malekova, L. et al. Inhibitory effect of DIDS, NPPB, and phloretin on intracellular chloride channels. Pflugers Arch. 455, 349–357 (2007).

76. Miller, C. & White, M. M. Dimeric structure of single chloride channels from Torpedo electroplax. Proc Natl Acad Sci USA 81, 2772–2775 (1984).

77. Matulef, K. et al. Discovery of potent CLC chloride channel inhibitors. ACS Chem. Biol. 3, 419–428 (2008).

78. Diaz, R. J. et al. Chloride channel inhibition blocks the protection of ischemic preconditioning and hypo-osmotic stress in rabbit ventricular myocardium. Circ. Res. 84, 763–775 (1999).

79. Diaz, R. J. et al. Enhanced cell-volume regulation in cyclosporin A cardioprotection. Cardiovasc. Res. 98, 411–419 (2013).

80. Tulk, B. M., Schlesinger, P. H., Kapadia, S. A. & Edwards, J. C. CLIC-1 Functions as a Chloride Channel When Expressed and Purified from Bacteria. Journal of Biological Chemistry 275, 26986–26993 (2000).

81. Guinamard, R., Simard, C. & Del Negro, C. Flufenamic acid as an ion channel modulator. Pharmacol. Ther. 138, 272–284 (2013).

82. Lama, L. et al. Development of human cGAS-specific small-molecule inhibitors for repression of dsDNA-triggered interferon expression. Nat. Commun. 10, 2261 (2019).

83. Vincent, J. et al. Small molecule inhibition of cGAS reduces interferon expression in primary macrophages from autoimmune mice. Nat. Commun. 8, 750 (2017).

84. Wang, M., Sooreshjani, M. A., Mikek, C., Opoku-Temeng, C. & Sintim, H. O. Suramin potently inhibits cGAMP synthase, cGAS, in THP1 cells to modulate IFN-β levels. Future Med. Chem. 10, 1301–1317 (2018).

85. Ni, G., Konno, H. & Barber, G. N. Ubiquitination of STING at lysine 224 controls IRF3 activation. Sci. Immunol. 2, (2017).

86. Wan, W. et al. STING directly recruits WIPI2 for autophagosome formation during STING-induced autophagy. EMBO J. 42, e112387 (2023).

87. Liu, D. et al. STING directly activates autophagy to tune the innate immune response. Cell Death Differ. 26, 1735–1749 (2019).

88. Prabakaran, T. et al. Attenuation of cGAS-STING signaling is mediated by a p62/SQSTM1-dependent autophagy pathway activated by TBK1. EMBO J. 37, (2018).

89. Wachter, R. M. & Remington, S. J. Sensitivity of the yellow variant of green fluorescent protein to halides and nitrate. Curr. Biol. 9, R628–9 (1999).

90. Kaizuka, T. et al. An Autophagic Flux Probe that Releases an Internal Control. Mol. Cell 64, 835–849 (2016).

91. Zhang, K., Wang, S., Gou, H., Zhang, J. & Li, C. Crosstalk Between Autophagy and the cGAS-STING Signaling Pathway in Type I Interferon Production. Front. Cell Dev. Biol. 9, 748485 (2021).

92. Wang, C. et al. Manganese Increases the Sensitivity of the cGAS-STING Pathway for Double-Stranded DNA and Is Required for the Host Defense against DNA Viruses. Immunity 48, 675-687.e7 (2018).

93. Lv, M. et al. Manganese is critical for antitumor immune responses via cGAS-STING and improves the efficacy of clinical immunotherapy. Cell Res. 30, 966–979 (2020).

94. Zhou, C. et al. Transfer of cGAMP into Bystander Cells via LRRC8 Volume-Regulated Anion Channels Augments STING-Mediated Interferon Responses and Anti-viral Immunity. Immunity 52, 767-781.e6 (2020).

95. Chu, T.-T. et al. Tonic prime-boost of STING signalling mediates Niemann-Pick disease type C. Nature 596, 570–575 (2021).

96. Gui, X. et al. Autophagy induction via STING trafficking is a primordial function of the cGAS pathway. Nature 567, 262–266 (2019).

97. Zhang, Q., Li, Y., Jian, Y., Li, M. & Wang, X. Lysosomal chloride transporter CLH-6 protects lysosome membrane integrity via cathepsin activation. J. Cell Biol. 222, (2023).

98. Feng, X., Liu, S. & Xu, H. Not just protons: Chloride also activates lysosomal acidic hydrolases. J. Cell Biol. 222, (2023).

99. Kellokumpu, S. Golgi ph, ion and redox homeostasis: how much do they really matter? Front. Cell Dev. Biol. 7, 93 (2019).

100. Kaunitz, J. D., Gunther, R. D. & Sachs, G. Characterization of an electrogenic ATP and chloride-dependent proton translocating pump from rat renal medulla. J. Biol. Chem. 260, 11567–11573 (1985).

101. Sepela, R. J. & Sack, J. T. Taming unruly chloride channel inhibitors with rational design. Proc Natl Acad Sci USA 115, 5311–5313 (2018).

102. Saha, S., Prakash, V., Halder, S., Chakraborty, K. & Krishnan, Y. A pH-independent DNA nanodevice for quantifying chloride transport in organelles of living cells. Nat. Nanotechnol. 10, 645–651 (2015).

