## Supplementary material for "Chloride Homeostasis Regulates cGAS-STING Signaling"

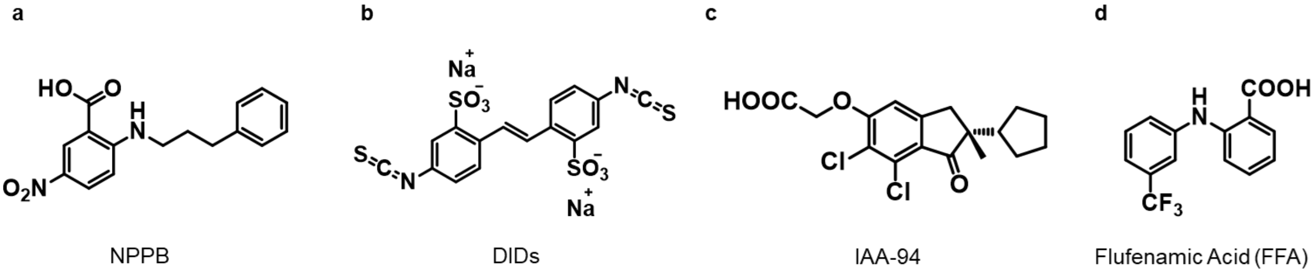


**Supplementary Figure 1.** Chemical structure of chloride channel blockers used in this study.

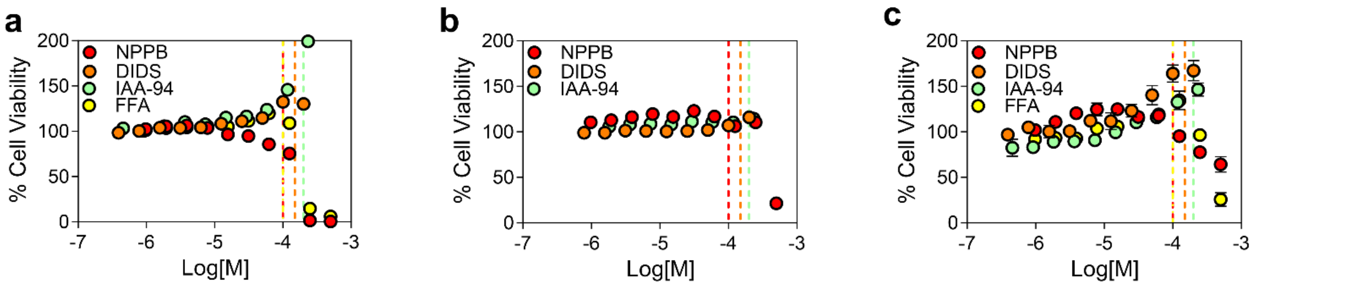


**Supplementary Figure 2.** Cell viability curve of (**a**) THP-1, (**b**) RAW 264.7 macrophage, and (**c**) primary human dermal fibroblast upon incubation of indicated chloride channel blockers overnight. Dashed lines indicate concentrations of compounds used throughout study. Error bars indicate the mean ± standard error of the mean (s.e.m.) of three independent measurements.

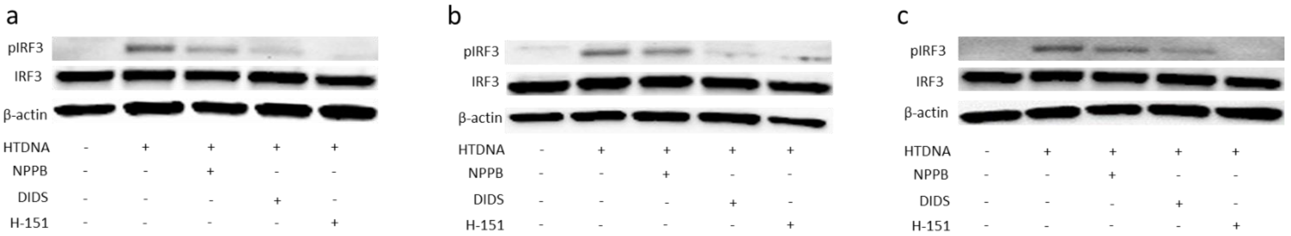


**Supplementary Figure 3.** Western blotting to measure the protein expression level of p-IRF3, IRF3 and β-actin in THP-1 cells that were pretreated with 100 μM NPPB,150 μM DIDS, and 15 μM H-151 overnight and then transfected with $2 \times 10^{-2}$ μg/μL HT-DNA overnight. Experiments were performed in three biological replicates.

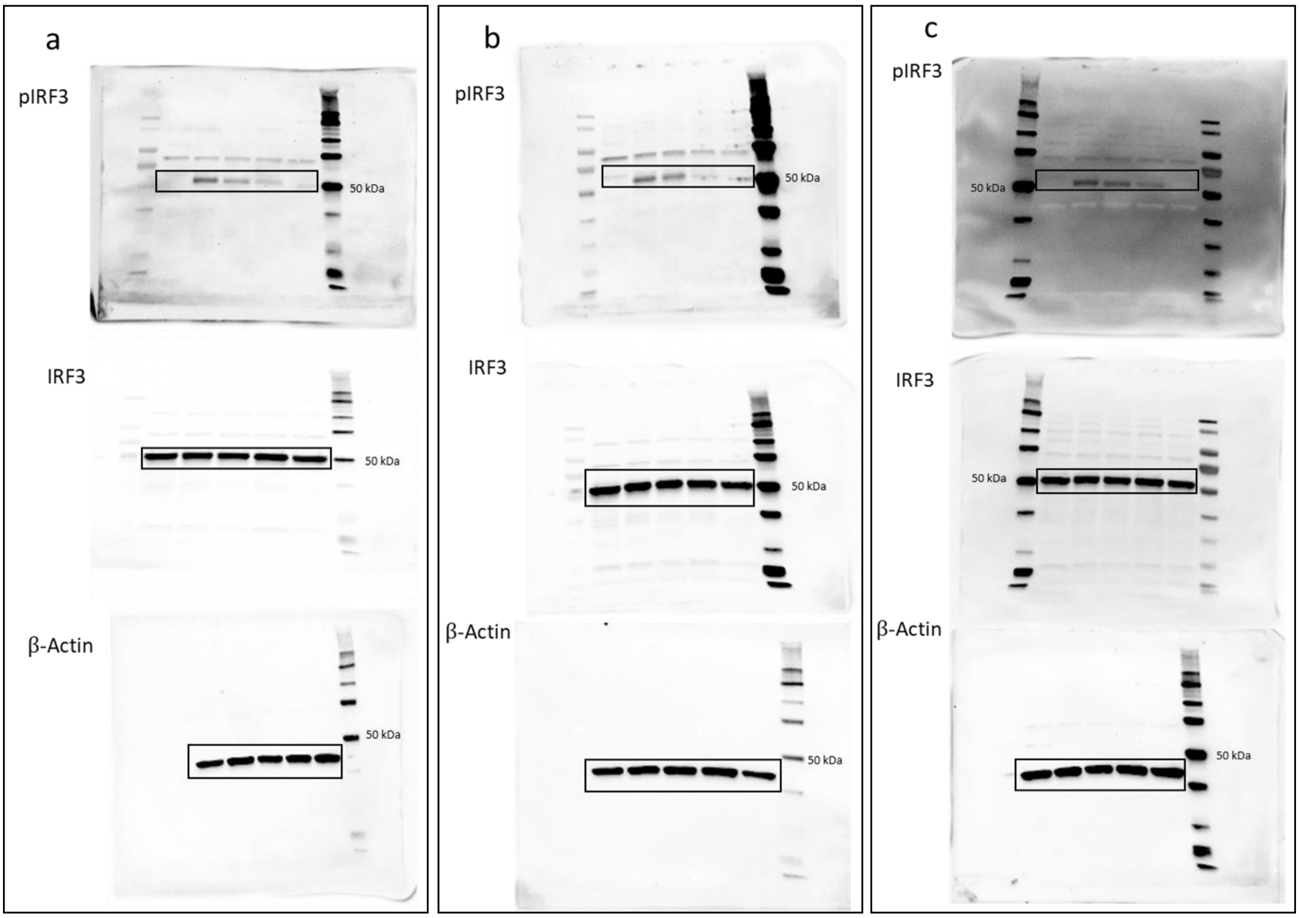


**Supplementary Figure 4.** Source data of Figure 1i and supplementary Figure 3.

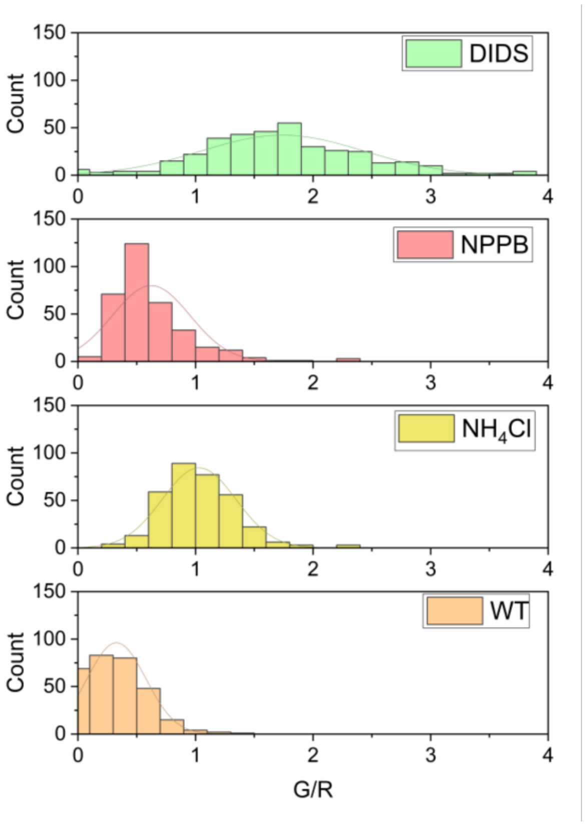


**Supplementary Figure 5.** Histogram analysis of the G/R values correlating to Figure 2(f−g) (>25 cells and >100 lysosomes per trial). Experiments were performed in triplicate.

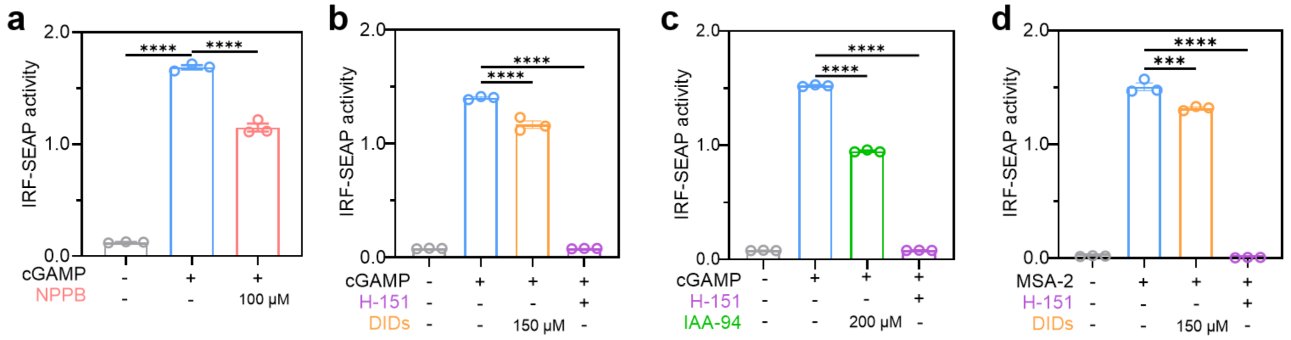


**Supplementary Figure 6.** THP1-Blue ISG cells were pretreated with (**a**) 100 μM NPPB, (**b**) 150 μM DIDS, (**c**) 200 μM IAA-94, and 15 μM H-151 for 1 h and then stimulated with 100 μM 2,3-cGAMP overnight. (**d**) THP1-Blue ISG cells were pretreated with 150 μM DIDS and 15 μM H-151 for 1 h and then stimulated with 30 μM MSA-2 overnight. Error bars indicate the mean ± standard error of the mean (s.e.m.) of three independent measurements. ****$P < 0.0001$. One-way analysis of variance (ANOVA) followed by Dunnett's test for multiple comparison.

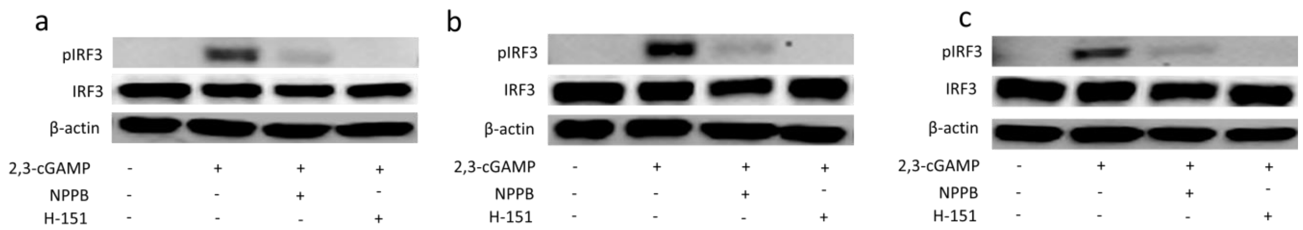


**Supplementary Figure 7.** Western blotting to measure the protein expression level of p-IRF3, IRF3 and β-actin in THP-1 cells that were pretreated with 100 μM NPPB, and 15 μM H-151 overnight and then stimulated with 100 μM 2,3-cGAMP overnight. Experiments were performed in three biological replicates.

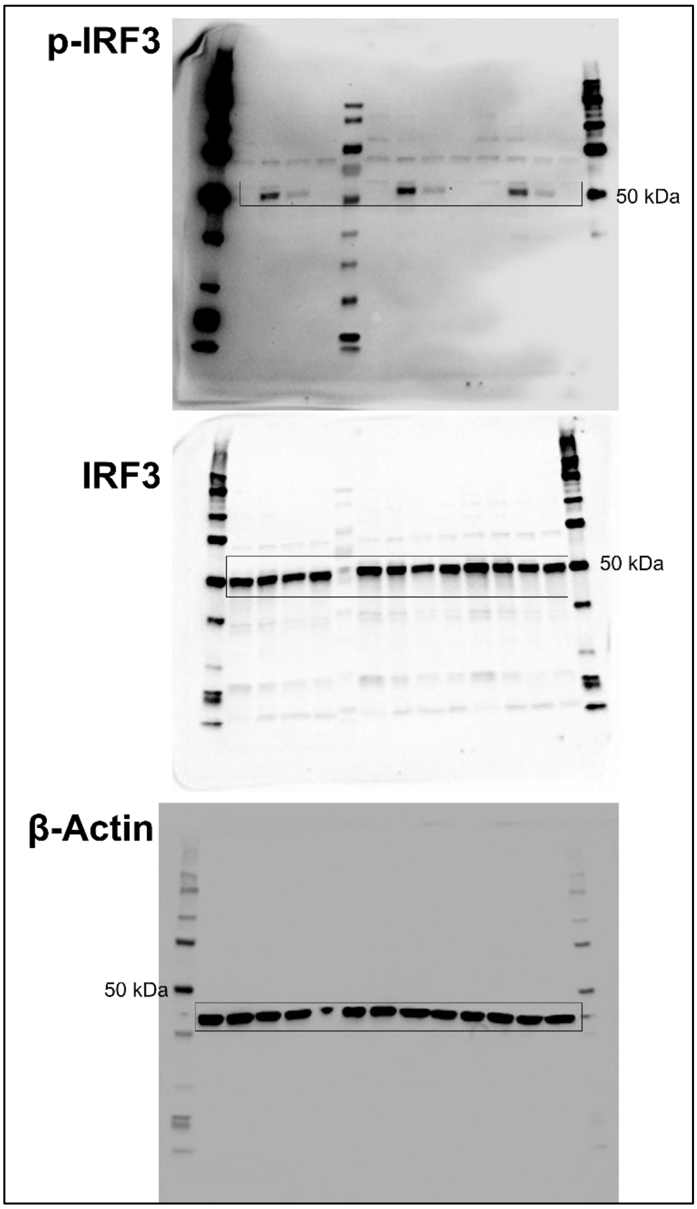


**Supplementary Figure 8.** Source data of Figure 4f and supplementary Figure 7.

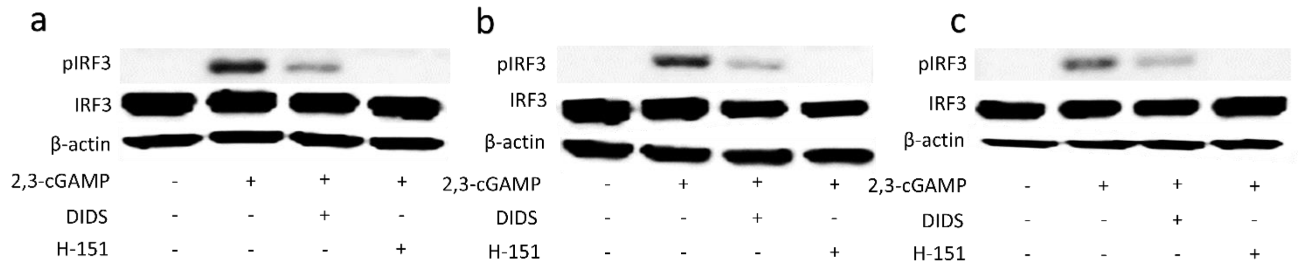


**Supplementary Figure 9.** Western blotting to measure the protein expression level of p-IRF3, IRF3 and β-actin in THP-1 cells that were pretreated with 150 μM DIDS, and 15 μM H-151 overnight and then stimulated with 100 μM 2,3-cGAMP overnight. Experiments were performed in three biological replicates.

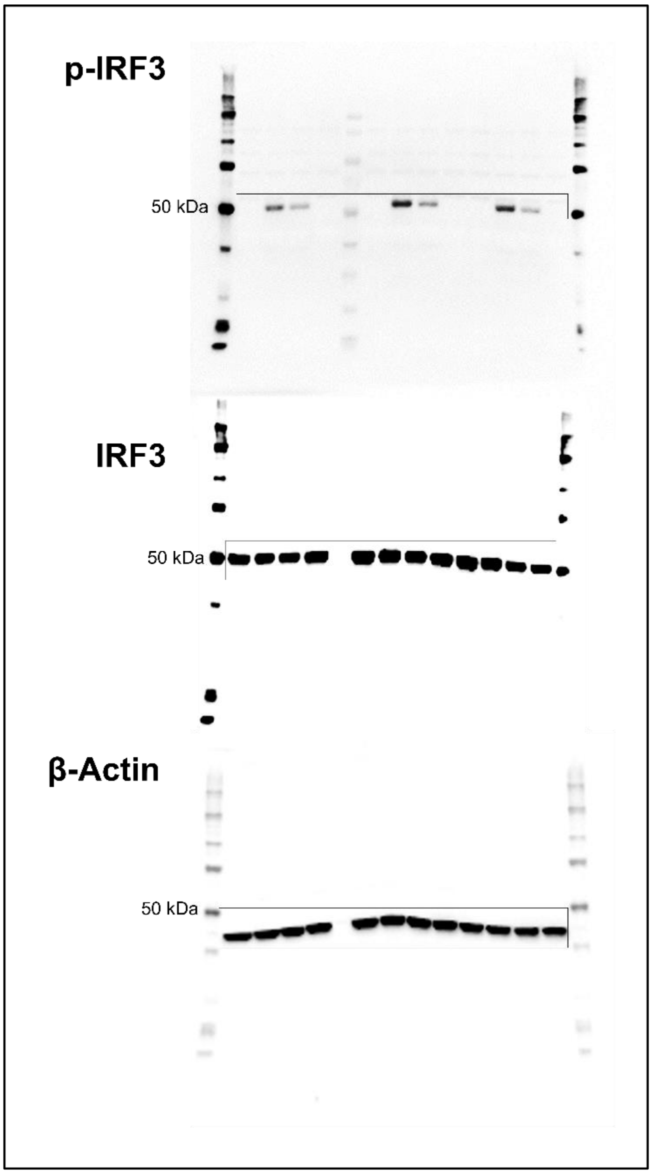


**Supplementary Figure 10.** Source data of Figure 4g and supplementary Figure 9.

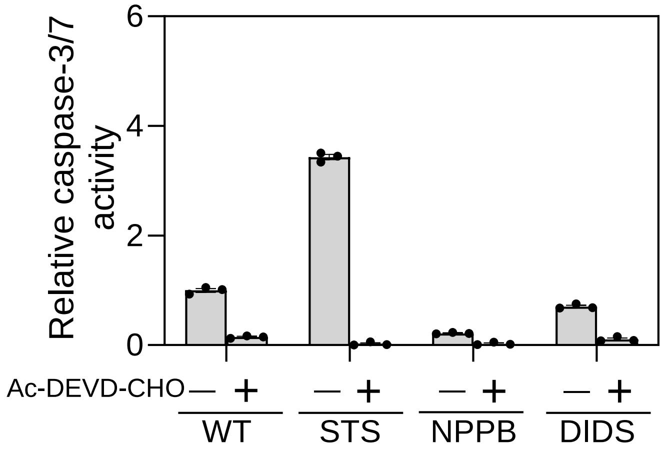


**Supplementary Figure 11.** Caspase-3/7 activity in RAW 264.7 macrophage. Cells were pretreated with 2 µM staurosporine (STS), 100 µM NPPB, and 150 µM DIDS overnight. 20 µM Ac-DEVD-CHO (caspase inhibitor) was added before analysis as a negative control. Error bars indicate the mean ± standard error of the mean (s.e.m.) of three independent measurements.

**Supplementary Table 1:** Information of cells used in this study.

| Sample | Sample Number | Gene | Gene Mutation | Age (at sampling) |
|---|---|---|---|---|
| NPC1-P1 | GM17913 | NPC1 | VAL1165MET | M (23 YR) |
| NPC1-P2 | GM18388 | NPC1 | ARG404GLN | F (No Data) |
| NI-1 | GM22277 | N/A | N/A | M (1 DA) |
| NI-2 | HDF (ATCC) | N/A | N/A | No Data |

DA = day, Yr = year.

**Supplementary Table 2:** Information of primers used in this study.

| Name | Sequence (5' to 3') | Length |
|---|---|---|
| Human GAPDH-F | ATCACCATCTTCCAGGAGCG | 20 |
| Human GAPDH-R | GGGCAGAGATGATGACCCTT | 20 |
| Human IFN-ß-F | AGGACAGGATGAACTTTGAC | 20 |
| Human IFN-ß-R | TGATAGACATTAGCCAGGAG | 20 |
